# Peptide:MHC Binding Stability Prediction Using Protein Language Models

**DOI:** 10.64898/2026.06.28.735023

**Authors:** Dhuvarakesh Karthikeyan, Benjamin Vincent, Alexander Rubinsteyn

## Abstract

Peptide:MHC class I (pMHC-I) binding stability governs the persistence of antigenic complexes at the cell surface and plays a key role in facilitating downstream immunological signals such as antigen presentation, T-cell activation, and immunodominance. However, methods for *in silico* stability prediction remain underexplored relative to binding affinity prediction, in part because available half-life datasets are sparse and expensive to collect. Here, we perform a systematic reassessment of pMHC-I stability prediction using controlled, similarity-aware data splits and apply a recently introduced supervised transfer-learning strategy to MINT, an interaction-aware protein language model, pre-trained on binding affinity and fine-tuned for quantitative half-life prediction. We show that MINT improves stability prediction over standard ESM-2 representations and existing predictors, and that assay-conditioned recalibration corrects systematic shifts across experimental measurement modalities. Across eluted ligand, immunogenicity, and personalized neoantigen prioritization benchmarks, predicted stability provides signal beyond binding affinity, enriching for naturally presented and immunogenic peptides within affinity-filtered candidate sets. These results establish pMHC-I half-life as an orthogonal and transferable biophysical signal connecting peptide binding, surface presentation, and T-cell recognition, and provide a leakage-aware, assay-aware framework for future antigen-presentation modeling.

## 1 Introduction

Immunogenicity, or the propensity of certain antigens to elicit immune responses in immunocompetent hosts, is a central component of the immune response and drives outcomes in antigen-directed therapies, including viral and neoantigen (NeoAg) vaccines as well as undesired anti-drug immune responses (ADIRs) against protein-based therapeutics. For T cells, whose evolutionary niche involves maintaining intracellular immunity by interfacing with cytosolic protein fragments presented on the surface of most cells, a successful immune response is predicated on concerted events in both the antigen-presenting cell (APC) and its engaging T cell. Core to antigen presentation, highly diverse yet specialized molecules known as major histocompatibility complex (MHC-I and MHC-II) molecules bind peptides from homeostatic protein turnover and present them at the cell surface for recognition by cytotoxic (CD8+) or helper (CD4+) T cells, respectively, enabling the detection of anomalous peptide signatures such as those associated with viral infection or malignancy. Of the many steps in the antigen presentation pathway: proteasomal cleavage, TAP-mediated transport, and competitive binding to available MHC-I molecules, peptide binding to MHC-I is widely considered the rate limiting step [1, 2] in determining the MHC ligandome, and has accordingly been the focus of computational efforts to model antigen presentation.

Presentation of a stable pMHC affects the immunogenicity of a peptide through two distinct but interrelated biophysical quantities: binding affinity, reflecting the thermodynamic favorability of the peptide:MHC association, and binding stability, reflecting the kinetic half-life of the resulting pMHC complex at the cell surface. While some basal level of both is required to mount an effective response, the two can play compensatory roles. For example, a peptide may bind the MHC-I groove with high affinity, yet form a complex that dissociates rapidly at the cell surface, negating its ability to stimulate a T cell. High peptide abundance can partially compensate for rapid dissociation by continually replenishing surface pMHC complexes, whereas a low-abundance peptide may still generate detectable antigenic signal if its pMHC complex is sufficiently long-lived. Conversely, a low abundance peptide with a high persistence may remain presented long enough to stimulate a cognate T cell. Of consequence, multiple studies have reported pMHC-I complex stability correlates more strongly with immunogenicity than binding affinity [3, 4], and more recently, that stability better predicts immunodominance among competing epitopes [5].

Despite this emerging significance, binding stability prediction remains far less supported than binding affinity, for which decades of sustained effort have produced large-scale experimental datasets and a mature ecosystem of accurate computational tools [6–8]. The experimental challenges associated with measuring complex dissociation at scale, including historically low-throughput methods that must account for inherently longer half-lives for more stable complexes, have resulted in comparatively sparse stability datasets and cover far fewer peptide-allele combinations. Only two maintained publicly available stability predictors exist: NetMHCstabpan [9], successor to NetMHCstab [3] and the first pan-specific neural network-based model, and TLStab [10], which leverages transfer learning from affinity and eluted ligand pre-training to improve binding stability prediction. While these methods represent important progress, their evaluation has been constrained by the scarcity, redundancy, and heterogeneity of available stability data, making published performance estimates difficult to compare directly and potentially optimistic due to uncontrolled data leakage, both within-task (high sequence similarity between training and evaluation peptides) and cross-task (overlap between binding affinity training sets and stability evaluation sets), that may inflate apparent generalization. Moreover, prior evaluations have largely emphasized rank-based metrics, leaving the accuracy of absolute half-life recovery under-quantified.

In this work, we systematically examine available pMHC-I binding stability data, characterizing sources of redundancy, noise, assay heterogeneity, and sequence-level leakage that can inflate reported model performance. We reevaluate existing predictors under stringent, similarity-aware data splits and apply the MINT architecture [11], an ESM-2 [12] variant explicitly pre-trained on protein–protein interactions and previously validated for binding affinity prediction, to the task of quantitative pMHC-I half-life prediction. To our knowledge, this represents the first application of a protein language model to pMHC-I binding stability prediction. Motivated by discordance in measured stability across assay types, we further introduce explicit assay-specific conditioning, showing that modeling experimental context yields stability estimates better aligned with distinct measurement modalities. We then ask whether these refined stability estimates transfer along the biological cascade from peptide:MHC binding and complex persistence to surface presentation, immunogenic recognition, and personalized neoantigen prioritization. Across curated population-level eluted ligand and immunogenicity datasets, as well as four clinical neoantigen cohorts, predicted pMHC-I stability provides signal complementary to binding affinity and improves identification of immunogenic candidates where existing stability predictors often fail. Together, these analyses establish a leakage-aware and assay-aware framework for pMHC-I stability prediction and position pMHC half-life as a distinct, transferable biophysical signal linking antigen presentation, T-cell recognition, and personalized immunotherapy design.

## 2 Methodology

### 2.1 Problem Formulation

In this work, we frame pMHC-I stability prediction as a regression task: given an arbitrary peptide sequence and an associated HLA allele (human leukocyte antigen, the human MHC allele), predict the dissociation half-life *t*_1*/*2_ (hours) of the resulting complex. Unlike methods that operate on 34-residue pseudo-sequences [13], we condition on the complete MHC-I protein sequence, enabling not only structural context beyond the immediate peptide binding residues, which can be significant for entire complex stability [14–17], but also seamless incorporation of HLA mutants as well. This full-sequence formulation thus provides a single, unified framework applicable to any MHC-I allele for which a protein sequence is available, including rare or uncharacterized variants absent from pseudo-sequence dictionaries.

To account for the scarcity of available binding stability data relative to binding affinity, we adopt two major design choices for our experiments. The first, borrowed directly from TLStab [10], which used the same principle in the same context, treats binding affinity prediction as a supervised pre-training task, adapting the model’s representations to the pMHC domain before fine-tuning on stability. The second deals with the variance in the half-life values themselves, which are collected under different experimental assay protocols (scintillation proximity, fluorescence, thermostability) at varying incubation temperatures, producing measurements that are not directly comparable across conditions. We address this heterogeneity problem by, for the first time in this setting, explicitly conditioning on assay type and temperature rather than treating them as unmodeled confounding variables:

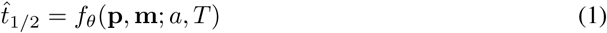

Where

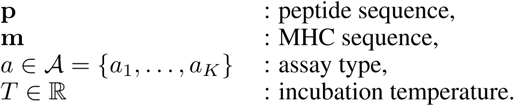

### 2.2 Datasets

A central concern in evaluating machine learning models in biology is that of data leakage: peptides in the evaluation set that share high sequence similarity with training peptides can inflate apparent performance. This phenomenon can become more obfuscated when multiple tasks are introduced, and yet its significance and ability to poison validation sets remain. To promote a faithful representation of our models’ performance, we adopt distinct splitting strategies for the BA and BS datasets, reflecting their different roles in our training pipeline. We assembled datasets spanning the full modeling pipeline: binding affinity pre-training, binding stability fine-tuning, assay-conditioned recalibration, and downstream functional evaluation. Full curation, allele normalization, decontamination, and split construction procedures are provided in *Supplementary Information (SI)*.

#### Binding affinity data

Binding affinity (BA) data were obtained from the NetMHCpan 4.1 training corpus [6]. Labels were represented using the standard transformed affinity score,

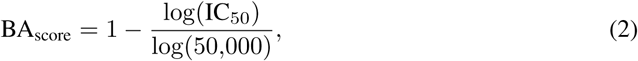

where larger values indicate stronger binding. Alleles were normalized to two-field HLA nomenclature and mapped to full-length MHC-I heavy chain sequences using IPD-IMGT/HLA [18]. After removal of non-human alleles and unmappable HLA sequences, the final BA dataset contained 170,107 measurements across 109 HLA-I alleles. Because BA data were used only for supervised pre-training, we applied an 80/10/10 split followed by peptide-level deduplication to eliminate exact peptide overlap between training and evaluation partitions, yielding 126,683 training, 21,712 validation, and 21,712 test examples.

#### Binding stability data

Binding stability (BS) measurements were obtained from the NetMHCstabpan training corpus [9], consisting of 9-mer peptide::MHC-I complexes with experimentally measured half-lives from scintillation proximity assays. After allele normalization and MHC sequence mapping, the dataset contained 27,034 measurements across 72 HLA-I alleles. Stability labels were modeled as log-transformed half-lives,

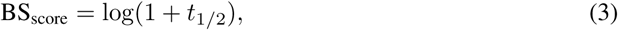

which preserves dynamic range across long-lived complexes and is mapped back to hours at inference by *t̂*_1*/*2_ = exp(*ŷ*) − 1. To reduce sequence-level leakage, unique peptides were clustered at an 80% sequence-identity threshold using normalized Levenshtein distance, and entire clusters were assigned to train, validation, or test partitions. This produced 21,626 training, 2,705 validation, and 2,700 test examples, with no evaluation peptide sharing ≥80% sequence identity with any training peptide in either the binding stability or binding affinity training partitions.

#### Independent IEDB stability holdout

To evaluate transfer beyond the NetMHCstabpan distribution, we constructed an independent holdout from IEDB MHC-I binding stability measurements [19]. Records were filtered to valid 8-15mer peptides with standard amino acids, quantitative half-life measurements, mappable HLA-I alleles, and full-length MHC-I sequences. Half-lives were converted to hours, and duplicate peptide:MHC measurements were aggregated by median half-life. To prevent both within-task and cross-task leakage, peptides sharing ≥80% sequence identity with any peptide used in the BA or BS training pipeline were removed. The final IEDB holdout contained 1,130 rows of unique, median-aggregated, pMHC-I complexes across 27 HLA-I alleles.

#### Assay-conditioned stability data

For assay-conditioned recalibration, we returned to the full IEDB stability export and retained quantitative MHC-I half-life measurements passing the same peptide, allele, and sequence-mapping filters. Measurements were assigned to four assay groups based on IEDB assay metadata: scintillation proximity assay (SPA), purified fluorescence (PF), cellular fluorescence (CF), and other. Incubation temperatures were extracted from assay comments when available and otherwise assigned protocol-specific defaults. Replicate measurements for the same peptide–allele–assay–temperature combination were aggregated by median half-life. After peptide-level decontamination against the held-out test set and iterative removal of any similar validation peptides, the final Stage 3 dataset comprised 8,465 training, 543 validation, and 1,133 held-out test entries (reclaiming three instances where peptides were measured using multiple assays, compared to the previous section’s *n*=1,130), with no peptide:MHC sequence overlap between splits and a clean less than 80% sequence identity to any training partition: binding affinity (Stage 1), binding stability (Stage 2), or assay-conditioning (Stage 3).

#### Downstream functional evaluation

To test whether predicted stability transfers to biologically downstream readouts, we evaluated all models on eluted ligand (EL), immunogenicity (IMM), and personalized neoantigen therapeutics benchmarks. No EL or immunogenicity labels were used during model training. EL data were drawn from the NetMHCpan 4.1 ligand corpus [6], filtered to human HLA-I alleles, normalized to two-field HLA nomenclature, and mapped to full-length MHC sequences. This dataset was then downsampled using stratified subsampling to one million peptide-allele pairs while preserving the original positive rate of approximately 5.4%. Immunogenicity data were obtained from IEDB T-cell assays [19] (March 2026 export), filtered to CD8^+^ responses with valid 8-15mer peptides and mappable HLA-I restrictions, and aggregated by unique peptide-allele pairs with binary labels indicating whether any positive T-cell response was reported. Finally, we curated four personalized neoantigen immunogenicity cohorts spanning multiple therapeutic modalities: NeoVax [20, 21] and PCV [22] synthetic long peptide vaccines, iNeST [23, 24] an mRNA vaccine, and NeoStim [25] an adoptive T-cell therapy. These cohorts were harmonized into a common patient-level schema containing mutant peptide sequences, HLA-I alleles, and binary immunogenicity labels, enabling per-patient evaluation of whether predicted stability provides signal complementary to binding affinity for neoantigen prioritization in an individualized, clinically relevant setting.

### 2.3 Model Architecture

#### 2.3.1 Tokenization and Input Representation

For tokenizing multimers, MINT [11] introduces a useful chain_id masking scheme that enables computation of the entire multimer with a single forward pass, which we retain and extend to our ESM baselines. For peptide and MHC-I sequences, we tokenize them and add the special `<cls>` and `<eos>` tokens, padding each to the peptide and MHC maximum sequence length. The two are then concatenated to form a single input tensor:

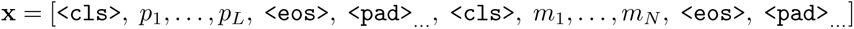

with its corresponding chain_id tensor:

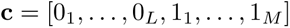

where *p*_1_*, . . ., p_l_* and *m*_1_*, . . ., m_n_* denote the peptide and MHC residue tokens, of length l and p, respectively padded out to length L and length N. No explicit separator token is inserted between the two chains. A parallel chain-ID tensor assigns identifier 0 to all peptide positions and identifier 1 to all MHC positions (including their respective special tokens and padding).

#### 2.3.2 Model

##### ESM-2

ESM-2 [12] is a protein language model pre-trained via masked language modeling on UniRef50 sequences [26]. We use the 650M-parameter variant (33 transformer layers, *d*=1,280), which learns residue-level representations that implicitly encode protein structure, recover inter-residue contacts from attention maps, and capture evolutionary fitness landscapes without structural supervision [27–29]. These properties have shown promise for peptide:MHC binding affinity prediction [30] but have not yet been evaluated in the binding stability setting.

##### MINT

MINT [11] extends ESM-2 650M with a dual-attention mechanism for modeling protein-protein interactions. Each transformer layer contains two independent multi-head attention modules with separate **Q**, **K**, **V** projections: one for intra-chain and one for cross-chain token pairs, distinguished by the chain-ID tensor. Intra- and cross-chain logits are assembled before a single softmax, and value aggregation routes each position through the corresponding module’s projections, doubling the attention parameters per layer. MINT was pre-trained on ∼96M protein-protein interactions from the STRING database [31], learning to distinguish intra-from inter-chain co-evolutionary signals via chain-aware masking. Additional architectural details and equations are provided in the *Supplementary Information (SI)*.

##### Downstream head

For the downstream stability prediction task, we append a two-layer projection head to the MINT/ESM backbone. After the final transformer layer, `<cls>`, `<eos>`, and `<pad>` tokens are masked out, and the remaining residue-level representations are mean-pooled across both chains to produce a fixed-dimensional embedding (**h**_pool_ ∈ ℝ^1280^). The projection head computes:

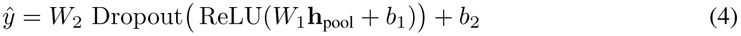

where 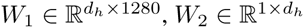, and *d_h_* is the projection hidden dimension.

##### Assay conditioning heads

To address systematic shifts in measured values depending on assay and temperature conditions, we introduce a lightweight conditioning module on top of the frozen models. Each training example is annotated with a categorical assay label *a* ∈ `{SPA, Purified_Fluor, Cellular_Fluor, Other}` and a continuous incubation temperature *t* (in ^◦^C). These are encoded as a learned assay embedding 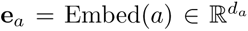 and a linear temperature projection 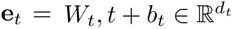.

Additional information on this aspect of our study may be found in the *Supplementary Information (SI)*. Briefly, we explored three conditioning architectures, all of which are initialized to recapitulate the trained predictions from the fine-tuning stage (identity initialization):

*(i) Additive fusion.* A residual vector is computed as **r** = MLP([**h**;, **e***_a_*;, **e***_t_*]) 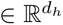 and added to the hidden state: *ŷ* = readout(**h** + **r**), again with zero-initialized output.
*(ii) Calibration.* The frozen Stage 2 scalar output *ŷ*^(*S*2)^ is affinely rescaled per assay (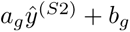, initialized to *a_g_* = 1*, b_g_* = 0) and corrected by a zero-initialized residual MLP: 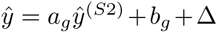, where Δ = MLP([**h**; **e***_a_*; **e***_t_*]).
*(iii) FiLM (Feature-wise Linear Modulation) [32].* The metadata and the Stage 2 hidden representation **h** are concatenated and passed through a two-layer MLP to produce sample-specific modulation parameters:

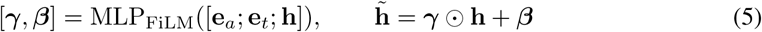

### 2.4 Model Training

#### 2.4.1 Transfer/Curriculum Learning Strategy

Building off of the success of transfer learning in TLStab [10], we similarly sought to use the relative abundance of binding affinity data compared to stability data to provide auxiliary training signal. Fasoulis et al. [10] demonstrated that transfer learning from binding affinity data substantially improves stability prediction. Motivated by these findings, we adopt a similar two-stage curriculum learning strategy using only binding affinity data for the pre-training stage, before extending this strategy to three-stage learning paradigm where the model is post-trained to provide assay-specific predictions:

##### Stage 1: Binding Affinity

Out of the box models (ESM-2 and MINT) are first tuned into pMHC representations by BA prediction using mean squared error (MSE) loss on the [0,1] BA-score labels. At this stage, the model sees most peptides and alleles, and learns preferences for sequence features when it comes to binding affinity.

##### Stage 2: Stability

Here, we take the model from Stage 1, including both the backbone weights and the projection head weights, and initialize Stage 2 training. In this stage, we freeze more of the backbone parameters, and lower the learning rate, allowing the affinity-tuned output mapping to serve as a prior for stability prediction. The model is then fine-tuned on the BS dataset with MSE loss on log(1 + *t*_1*/*2_)-transformed labels to reduce the influence of outliers with extreme half-lives.

##### Direct Fine-tuning Ablation

To quantify the contribution of the BA pre-training stage, we additionally train a hyperparameter-optimized model that skips Stage 1 and directly fine-tunes on the BS dataset from the pre-trained protein language models. We run a 2×2 factorial design of MINT vs. ESM-2 backbone crossed with two-stage vs. direct training to allow for the independent assessment of the contributions of interaction-aware pre-training and curriculum learning.

##### Stage 3: Assay Conditioning

Stage 2 is trained exclusively on SPA measurements at 37^◦^C derived from the NetMHCstabpan [9] training corpus. To extend predictions to the multi-assay landscape without overwriting the learned stability representations, we freeze the entire backbone and attach a lightweight conditioning module (∼810K trainable parameters) that modulates the Stage 2 hidden features based on a learned assay embedding and a continuous temperature encoding to shift and scale the predictions based on empirical assay-specific differences (the final model unfreezes the projection layer for marginal benefit; see *Supplementary Information (SI)*).

#### 2.4.2 Training Details

Stage 1 and Stage 2 models were optimized using AdamW [33] with *β* = (0.9, 0.98), *ɛ* = 10^−8^, and weight decay of 0.01. Gradients were clipped to a maximum norm of 1.0. A ReduceLROnPlateau scheduler halved the learning rate after 3 epochs without improvement in validation loss. Training was run for a fixed number of epochs (20 epochs for BA and 200 epochs for BS) and the checkpoint with the lowest validation loss was retained for downstream use. All experiments were conducted on 4× NVIDIA A6000 GPUs (48 GB each). Hyperparameter sweeps were parallelized across GPUs using Weights & Biases agents, with each agent assigned to a single GPU. Stage 3 switches to Huber loss (*δ* = 1.0) to preserve sensitivity in the smaller half life ranges while limiting the influence of extreme half-life outliers that arise when pooling across heterogeneous assay protocols.

##### Layer Freezing

To preserve pre-trained representations while allowing task-specific adaptation, we froze the bottom *k* of the 33 transformer layers, along with the token embedding and language modeling head parameters. The fraction of frozen layers was treated as a hyperparameter: we swept over freeze fractions of 0.3, 0.5, and 0.7 for Stage 1, and 0.5, 0.7, and 0.9 for Stage 2, reflecting the prior on the smaller stability dataset benefiting from greater regularization through freezing. The freeze rate used for each model was identified via sweep. Stage 3 freezes the backbone entirely (all 33 layers). In our benchmarks we freeze even the Stage 2 projection head, restricting gradient updates to the conditioning parameters alone. However, our production model trains the projection head as well, to marginal benefit. These reflect the design goals of recalibrating existing predictions for assay-specific perturbations of an intrinsic value.

##### Hyperparameter Selection

Hyperparameters were selected via Bayesian optimization sweeps executed via Weights & Biases [34], with validation loss as the objective. Sweeps were run for MINT’s two-stage training (individual sweeps for BA fine-tuning and BS fine-tuning), MINT direct fine-tuning, and each of the three Stage 3 conditioning architectures. While analogous sweeps were conducted using ESM-2, the optimal hyperparameters did not vary substantially from those found for MINT, and as such the MINT hyperparameters were applied to the ESM-2 baselines for a roughly parameter-equivalent comparison. For Stages 1 and 2, configs were run for 500 steps and the models with the lowest validation loss on the NetMHC partition were chosen; in the event of near ties, the configuration with the larger parameter count was selected, matched for learning rate. Stage 3 sweeps were run with the same protocol but with Huber or MSE loss as options. Table 1 summarizes the final hyperparameters for each model.

**Table 1:** Hyperparameters for ESM-2 and MINT stability prediction models.

| Parameter | ESM-2 |  | MINT (S1/S2) |  | Stage 3 |
| --- | --- | --- | --- | --- | --- |
|  | Direct | Transfer | Direct | Transfer | FiLM |
| Backbone | ESM2 | ESM2 | MINT | MINT | MINT |
| Cross-Attention | No | No | Yes | Yes | Yes |
| Learning Rate | $1 \cdot 10^{-4}$ | $1 \cdot 10^{-4} \rightarrow 3 \cdot 10^{-5}$ | $1 \cdot 10^{-4}$ | $3 \cdot 10^{-5} \rightarrow 3 \cdot 10^{-5}$ | $3 \cdot 10^{-5}$ |
| Hidden Dim | 256 | 512 | 256 | 512 | 512 |
| Dropout | 0.3 | $0.3 \rightarrow 0.2$ | 0.3 | $0.2 \rightarrow 0.2$ | 0.0 |
| Freeze % | 90% | $70\% \rightarrow 70\%$ | 90% | $50\% \rightarrow 70\%$ | $100\%^\dagger$ |
| Batch Size | 16 | 16 | 16 | 16 | 16 |
| Loss | MSE | MSE | MSE | MSE | Huber($\delta=1.0$) |
| Assay Emb Dim | — | — | — | — | 32 |
| Temp Emb Dim | — | — | — | — | 8 |
| Cond. Hidden Dim | — | — | — | — | 512 |
| Trainable Params | — | — | — | — | $\sim 810K^\dagger$ |

<sup>†</sup> The production S3 model unfreezes the Stage 2 projection head, increasing trainable parameters to $\sim 1.5M$ .

**Table 2:** Performance on the discovery (NetMHCstabpan) test set (*n* = 2,700).

| Method | $n$ | Spearman $\rho$ | Pearson $r$ | CCC | RMSE | log-RMSE | MAE |
| --- | --- | --- | --- | --- | --- | --- | --- |
| NetMHCstabpan<sup>†</sup> | 2,700 | 0.876 | 0.532 | 0.500 | 9.73 | 0.629 | 3.74 |
| TLStab<sup>‡</sup> | 2,700 | 0.698 | 0.306 | 0.132 | 45.55 | 0.929 | 8.91 |
| ESM-2 Direct | 2,700 | 0.574 | 0.515 | 0.340 | 9.61 | 0.869 | 4.33 |
| ESM-2 Transfer | 2,700 | <u>0.745</u> | <u>0.714</u> | **0.682** | <u>7.67</u> | <u>0.704</u> | <u>3.35</u> |
| MINT Direct | 2,700 | 0.732 | 0.679 | 0.637 | 8.04 | 0.735 | 3.65 |
| MINT Transfer | 2,700 | **0.791** | **0.761** | <u>0.673</u> | **7.33** | **0.650** | **3.22** |
| TLStab (filtered) | 378 | 0.461 | 0.313 | 0.301 | 9.98 | 0.985 | 4.62 |

<sup>†</sup> Trained on test evaluation data.
<sup>‡</sup> Trained on significant peptide overlap with evaluation data.

### 2.5 Baselines

To ensure a fair comparison across methods with different training corpora and architectural priors, we evaluate all models on two held-out sets: a *discovery* test partition drawn from the NetMHCstabpan corpus (filtered to *<*80% sequence identity against all BA and BS training peptides) and an *external* IEDB holdout similarly filtered to *<*80% sequence identity against the all training epitopes. Lastly, for all downstream functional evaluation studies, in order to retain statistical power, a looser peptide-level deduplication was used for each of the models’ upstream training data. Metrics are therefore reported on individualized subsets of the partition whose global numbers are shown wherever characterized. We compared the following methods under identical evaluation conditions:

- **NetMHCstabpan** [9]: a neural network ensemble model trained on the same NetMHCstabpan data distribution using pseudo-sequences. Because NetMHCstabpan was trained using CV and the production model was released having been trained on the entire dataset, its performance on the discovery test set represents an upper bound; performance on the IEDB holdout provides a fairer comparison.
- **TLStab** [10]: a transfer learning approach that pre-trains on binding affinity and eluted ligand data before fine-tuning for stability prediction, and immunogenicity prediction. TLStab uses a similar architecture to NetMHCstabpan and extends the training procedures.
- **ESM-2 Direct**: ESM-2 fine-tuned directly on BS data without Stage 1 pre-training.
- **ESM-2 Transfer**: the two-stage model with ESM-2 backbone.
- **MINT Direct**: MINT fine-tuned directly on BS data without Stage 1 pre-training.
- **MINT Transfer**: the full two-stage model (BA → BS) with MINT backbone.
- **NetMHCpan 4.1**^∗^ [6]: a pan-allele binding affinity predictor trained on binding affinity measurements and eluted ligand measurements by the same architecture. It is used only for the downstream predictions of immunogenicity and partially for the eluted ligand analysis.

NetMHCstabpan [9] and NetMHCpan 4.1 [6] predictions were obtained by running the publicly available executables. TLStab [10] predictions were obtained from running the published code accompanying the original study. All external predictions were converted from stability scores (*s*) to half-life in hours wherever applicable.

#### ESM-2 single-chain baseline

To isolate the contribution of cross-chain attention, we evaluate an ESM-2 baseline that shares the same backbone, tokenization, and projection head as MINT but replaces all cross-chain attention logits with −∞ masks, so that peptide and MHC tokens attend only within their own chain throughout all 33 layers. The two chains share no information at the representation level and are combined only at the final mean-pooling step. This is equivalent to two independent ESM-2 forward passes with a shared encoder, attributing any performance gap between MINT and ESM-2 specifically to the cross-chain attention mechanism.

### 2.6 Joint (BA + BS) Model Analysis

To evaluate whether stability predictions provide discriminative signal beyond binding affinity, we fit simple one or two feature logistic regression models (solver: L-BFGS, max iterations: 5,000) comparing a BA-only model (*X* = BA) against joint models (*X* = [BA, BS]), where BA denotes the predicted binding affinity value and BS denotes the predicted binding stability value. All stability values were transformed to (log(1 + *x*)) hours prior to fitting. All affinity values use the standard 1 − log_50_*_k_* transform. No additional feature scaling was applied. For eluted ligand prediction, models were evaluated on the full dataset and on subsets gated to the top 10%, 5%, and 2% of NetMHCpan 4.1 predicted binders; for immunogenicity, all samples were used without gating. Statistical significance was assessed via likelihood ratio tests comparing the nested BA-only and joint model log-likelihoods (LR = −2(*ℓ*_null_ − *ℓ*_alt_), *χ*^2^, *df* = 1), with adjusted *p*-values using the Benjamini-Hochberg procedure. Calibration improvement was quantified as the percent relative Brier skill score [35], computed across a 5-fold stratified cross-validation (stratified by label, seed = 42). Feature redundancy between each stability predictor and the BA feature was assessed using the Spearman-based variance inflation factor, 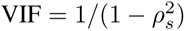, where *ρ_s_* is the Spearman rank correlation between the two features.

### 2.7 Metrics

Model performance and utility were assessed using metrics for both the continuous stability prediction as well as binary classification tasks at both the dataset-level (EL and IMM) and personalized antigen prioritization:

#### Continuous stability prediction

- **Spearman’s *ρ***: rank correlation between predicted and observed half-lives, capturing the model’s ability to correctly order peptides by stability. This serves as our primary metric, as it is invariant to monotonic transformations and robust to the nonlinear scaling differences that arise from heterogeneous experimental assays.
- **Pearson’s *r***: linear correlation coefficient, measuring the strength of the linear relationship between predictions and observations. Always computed in terms of hours.
- **log-RMSE**: 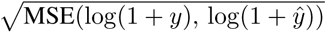, was used to attenuate the impact of extreme half-life values and better capture poor performance in the low-stability regime.
- **CCC** (Lin’s concordance correlation coefficient) [36]: 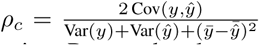, which jointly penalizes deviations in both correlation and calibration. Reported on log-values.

#### Binary classification

For eluted ligand (EL) and immunogenicity (IMM) evaluation, we report:

- **AUROC**: area under the receiver operating characteristic curve, measuring discrimination between positive and negative examples across all classification thresholds.
- **AUPRC**: area under the precision-recall curve, which is commonly reported under severe class imbalance, as it reflects a bias towards rightly prioritizing true positives.
- **Recall at 5% FPR**: sensitivity at a fixed 5% false-positive rate, capturing model performance at a high-specificity operating point relevant to experimental follow-up, where false positives carry substantial validation cost.
- **Precision@K and Recall @K**: The precision and recall computed on the top-*K* predicted values, taken by rank ordering the predictions and computing scores on the label prevalences. Useful in comparing performances in clinically relevant regimes.

## 3 Results

### 3.1 MINT outperforms existing stability predictors on NetMHCstabpan dataset

We first sought to evaluate the added benefit of transfer learning and the use of a multimer-aware protein language model over a standard pLM. In this preliminary analysis we evaluated four new pLM models (ESM Direct, ESM Transfer, MINT Direct, and MINT Transfer) against the established NetMHCstabpan [9] and TLStab [10] on the held-out test partition of the NetMHCstabpan data (*n* = 2,700) (**Figure 2a**), specifically curated to ensure zero overlap and leakage by sequence identity with the training sets of the upstream binding affinity and stability datasets (**Figure 2b; Supplementary Figure 2a-b**). While NetMHCstabpan achieved the highest Spearman correlation (*ρ* = 0.88), we find that among methods without data leakage, MINT Transfer achieved the best performance (Spearman *ρ* = 0.79, Pearson *r* = 0.76), with its Pearson *r* outperforming even NetMHCstabpan (**Figure 2c; Supplementary Figure 2c**). Of significance, we find that transfer learning helped both variants of ESM-2 and MINT, with improvements in Spearman *ρ* from 0.57 to 0.75 and 0.73 to 0.79, respectively. These were matched by improvements in Pearson *r* from 0.52 to 0.71 and 0.68 to 0.76 for ESM-2 and MINT as well, demonstrating that binding affinity representations transfer meaningfully to stability prediction. Notably, we find retaining the weights of the projection head from Stage 1 outperforms re-initialization, suggesting that the affinity-trained output mapping does provide a useful prior for the stability task. When controlling for transfer learning, we find that both variants of MINT outperformed ESM-2 (Direct vs Transfer), highlighting the added performance of using the multimer attention and pre-training on protein-protein interactions.

**Figure 1:**
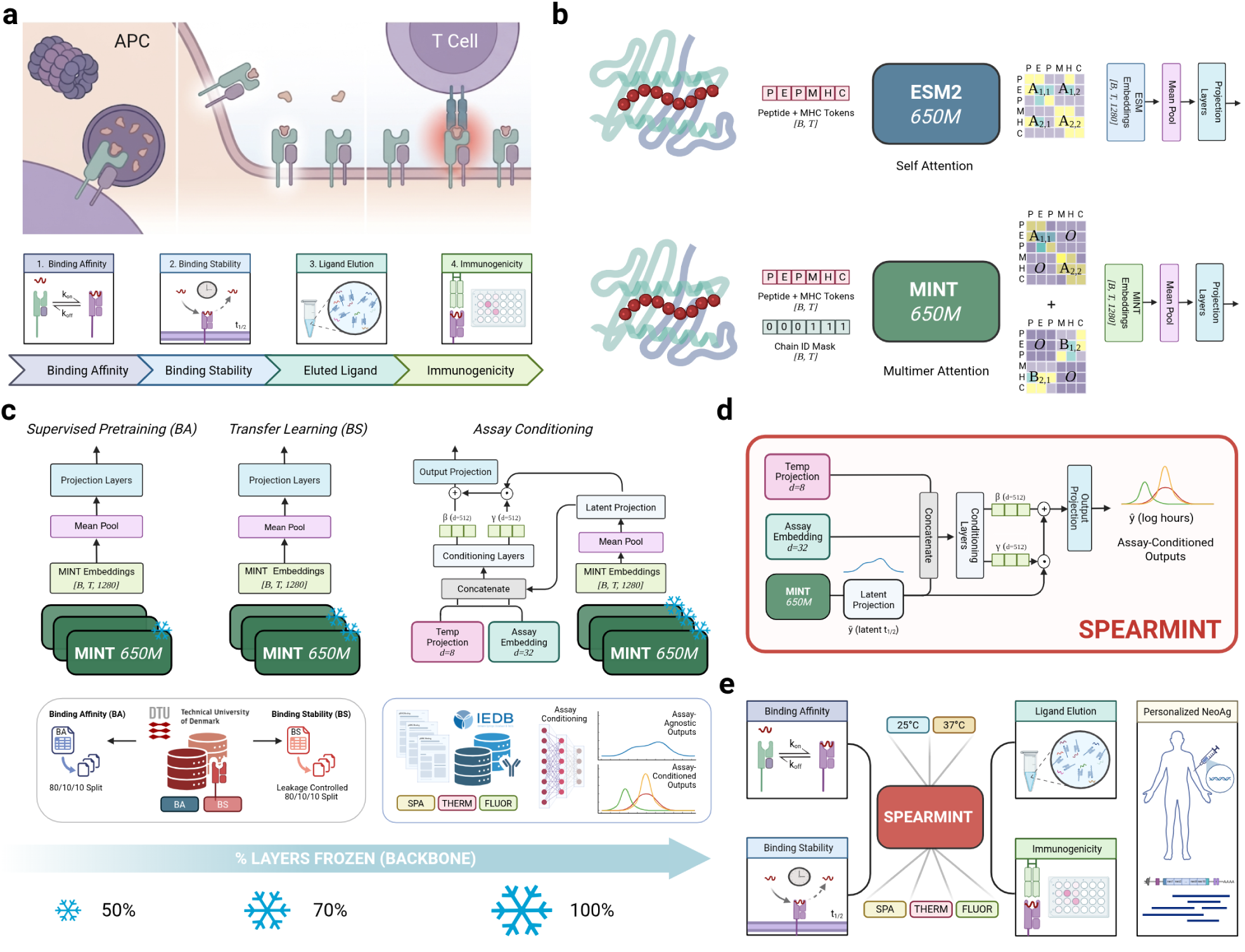
Overview of SPEARMINT. **a.** APC-centric determinants of immunogenicity, spanning proteasomal peptide generation, MHC-I loading competition, surface pMHC persistence, and downstream CD8+ T cell recognition. **b.** Comparison of the MINT architecture [11] and its progenitor, ESM2 [12], highlights the self-versus multimer-attention for capturing peptide:MHC interactions. **c.** Multi-stage training with progressive layer freezing from domain adaptive, supervised pre-training on binding affinity measurements, fine-tuning on binding stability measurements, and post-training recalibration of half-life values using assay-conditioning. **d.** Flagship SPEARMINT model architecture enables feature-wise modulation of learned stability signal to adapt to assay-level shifts in observed values. **e.** Robust benchmarks evaluate models’ ability to accurately predict observed half-life values along with evaluating stability models on downstream immune endpoints including eluted ligand presentation, immunogenicity, and neoantigen prioritization, assessing whether stability transfers to natural presentation and the subsequent propensity for T cell recognition.

**Figure 2:**
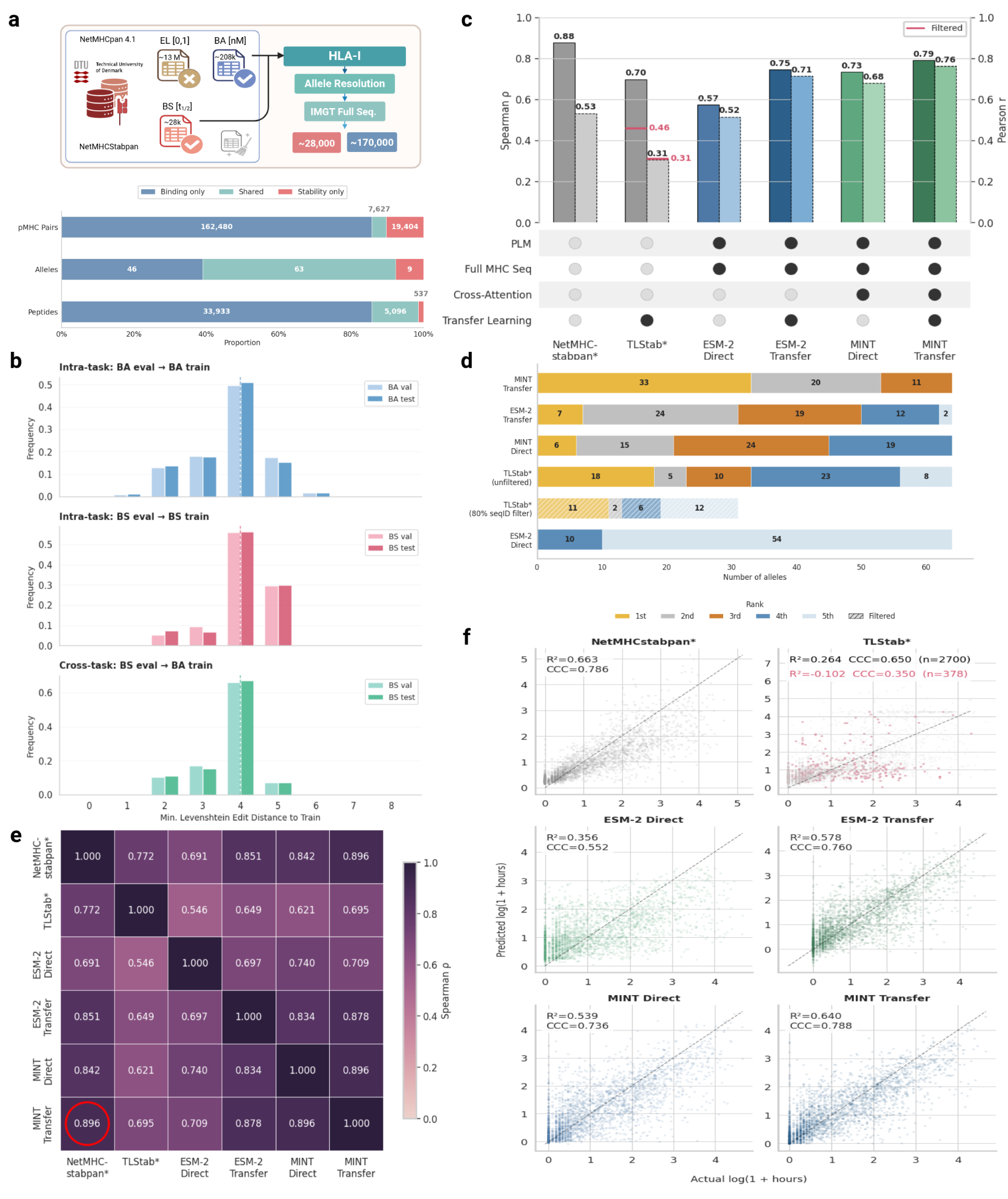
PPI-centric PLM and transfer learning yield more accurate predictions of pMHC half life on stringent holdout data. **a.** Multi-dataset creation and curation schematic diagram and accompanying inter-task overlap. **b.** Inter- and Intra-task edit distances between evaluation sets (validation, test) and train for binding affinity and binding stability datasets. **c.** Holdout validation results on NetMHCstabpan dataset. Spearman and Pearson correlation coefficient values are reported for all models. An asterisk is marked next to NetMHCstabpan and TLStab to indicate data leakage. A second set of values is reported for TLStab on a subset comprised of the data with an 80% sequence identity filter to their training set. **d.** Podium plot showing the per-allele rank of each model with 1st being best and 5th being worst. NetMHCstabpan excluded due to perfect leakage, TLStab shown as both filtered and unfiltered variants, with the other models being compared against TLStab unfiltered. **e.** Concordance analysis. NetMHCstabpan used as an oracle on this partition, with the model having the highest concordance to NetMHCstabpan highlighted in red. **f.** Correlation plot between predicted and actual log half-life values. *R*^2^ and Lin’s Concordance Correlation Coefficient (CCC) are reported per model.

To account for allelic heterogeneity in the dataset, and its clinically relevant repercussions, we evaluated our models’ performance on a more granular level, stratifying by presenting allele. When we analyzed ranks of Spearman values for each model across alleles, MINT Transfer and ESM-2 Transfer achieved the most ranks in the top three, closely followed by MINT Direct and then TLStab (**Figure 2d; Supplementary Figure 2d-e**). Of note, MINT Transfer was ranked first 33 times across alleles, the largest margin by any model. Using NetMHCstabpan as an oracle on this partition, we calculated the Spearman correlations between its predicted values and those of the remaining models. Here we found that MINT Transfer had the highest rank-agreement with NetMHCstabpan (*ρ* = 0.896), followed by ESM-2 Transfer (*ρ* = 0.851) and MINT Direct (*ρ* = 0.842), suggesting that the combination of structural pre-training and curriculum learning most closely recovers the ranking behavior of an oracle (**Figure 2e**). NetMHCstabpan is used as an oracle for this analysis because the final model was trained on this data and evaluated using cross-validation. Finally, in an effort to quantify the calibration of the models’ predictions, in terms of absolute error, we demonstrate MINT stability’s performant Lin’s concordance correlation coefficient (CCC) of 0.788, on par with NetMHCstabpan’s leakage inflated 0.786 (**Figure 2f**). In line with the other metrics, the second best model was ESM-2 Transfer followed by MINT Direct and ESM-2 Direct, with similar patterns in the errors (**Supplementary Figure 2f**).

### 3.2 All Stability Predictors Fare Worse on Heterogeneous IEDB Dataset

To assess generalization beyond the NetMHCstabpan distribution, comprised of data originating from a single group, we evaluated all methods on a curated subset of IEDB, ensuring controlled leakage to all upstream training sets (**Figure 3a**). Of particular interest, we identified early that pMHC stability for the same peptide-allele pair, measured using different assays and assay conditions not only gave different measurements, but also that the rank order of their measurements were discordant as well (**Figure 3b; Supplementary Figure 4a-c**). Of the most common assays, the Spearman correlation between SPA and purified fluorescence was 0.34, followed by 0.2 for SPA and cellular fluorescence. Though the number of data points were limited, we did find the highest inter-assay agreement of 0.72 between cellular and purified fluorescence, both of which used fluorescently labeled antibodies to measure dissociation. Given this, we analyzed the models stratified in accordance with the assay in which it was measured in. Interestingly, we found that all models’ performance fell below a Spearman *ρ* of 0.4, even for the SPA subset, highlighting a strong distribution shift between the training data and the data being generated and deposited to the IEDB (**Figure 3c**). While MINT Transfer led in the distribution on which the models were trained (SPA), it fell on both the purified and cellular fluorescence, underscoring the potentially protective capacity of smaller models as regularization. To investigate the confounds driving this change in performance we characterized the dataset in juxtaposition to the NetMHCstabpan data (**Supplementary Figure 3**). While we found that, in general, model performance degrades on alleles where the shift in label distribution is high (**Figure 3d; Supplementary Figure 4d**), the relationship was most explained by the fraction of examples per-allele that were measured using SPA, with a Spearman correlation between the performance and fraction SPA of 0.858 and 0.733 for the per-allele *ρ* and Pearson *r*, respectively (**Figure 3e-g; Supplementary Figure 4e-f**). We also demonstrate that this effect causes a drop in performance within allele as well, going from the subset that was measured using SPA versus the pMHCs measured using a different assay (**Figure 3h**).

**Figure 3:**
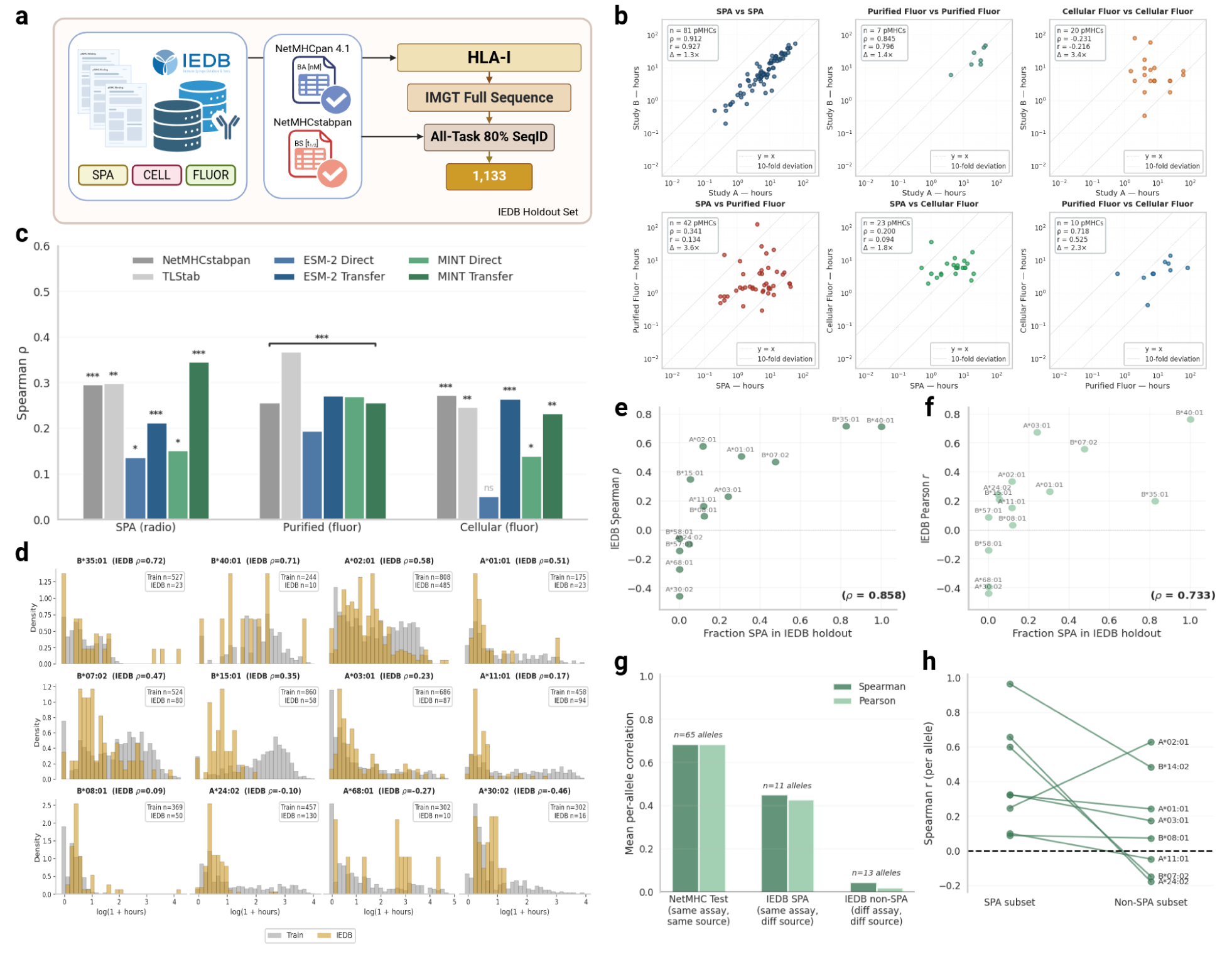
Heterogeneous IEDB assay conditions reveal assay-specific shifts in observed pMHC stability values that impact all stability prediction methods. **a.** IEDB heterogeneous assay generalization dataset construction diagram. **b.** Intra- and inter-assay consistency analysis for binding stabilities of the same pMHC measured using the same or different assays. Correlation statistics and median fold change between values (calculated as randomized Study B/Study A) are displayed. **c.** Assay-stratified performance benchmark. Significance levels are bootstrapped by computing the test statistic (Spearman *ρ*) on permuted labels. **d.** Analysis of distribution shift between training labels and held out IEDB test labels per allele. Distributions for the 12 most represented test alleles are shown with the corresponding MINT Transfer *ρ* values. **e-f.** Assay generalization analysis. MINT Transfer performance (Spearman and Pearson) per allele as a function of the fraction of data points derived by SPA (same assay as NetMHCstabpan training data). **g.** Bar plots of per-allele means of Spearman and Pearson correlation values for the NetMHCstabpan, IEDB SPA, and IEDB non-SPA evaluation set partitions for MINT Transfer. **h.** Paired dot plots show allele-level changes in MINT Transfer performance between held out examples in the IEDB given their assay of measurement (SPA vs. Non-SPA). Alleles with at least 5 samples per assay are shown.

### 3.3 Assay Conditioning

While the NetMHCstabpan SPA corpus provides a homogeneous historical foundation for pMHC-I stability modeling, the experimental landscape has since shifted away from radiolabeling toward thermostability readouts [37] and fluorescence-based half-life assays [38], limiting the field’s ability to generalize to newer data modalities. To improve our performance on non-SPA derived measurements, we investigated approaches to predict assay-specific half life predictions using the 8,465 multi-assay pMHC-I measurements (5,887 SPA, 2,468 purified fluorescence, 105 cellular fluorescence, and 5 other) from IEDB (**Figure 4a; Supplementary Figure 5**). Given the limited training data per assay, we adopt the strategy of conditioning the models’ outputs, as opposed to retraining the entire model, to learn the assay-specific shifts to a performant SPA prediction, reflecting the assumption that pMHC complexes possess an intrinsic binding stability that is systematically perturbed by assay-specific factors. We therefore introduced assay conditioning as an identity-initialized recalibration layer that preserves the shared stability estimate while learning protocol-specific corrections across SPA, purified fluorescence, and cellular fluorescence measurements. We benchmarked three different conditioning strategies of varying complexities, demonstrating feature-wise linear modulation (FiLM) [32] as the most performant (*Supplementary Information (SI)*). We henceforth refer to this approach and the resulting model as SPEARMINT (Stability Prediction of Epitopes with Assay Recalibration using MINT).

**Figure 4:**
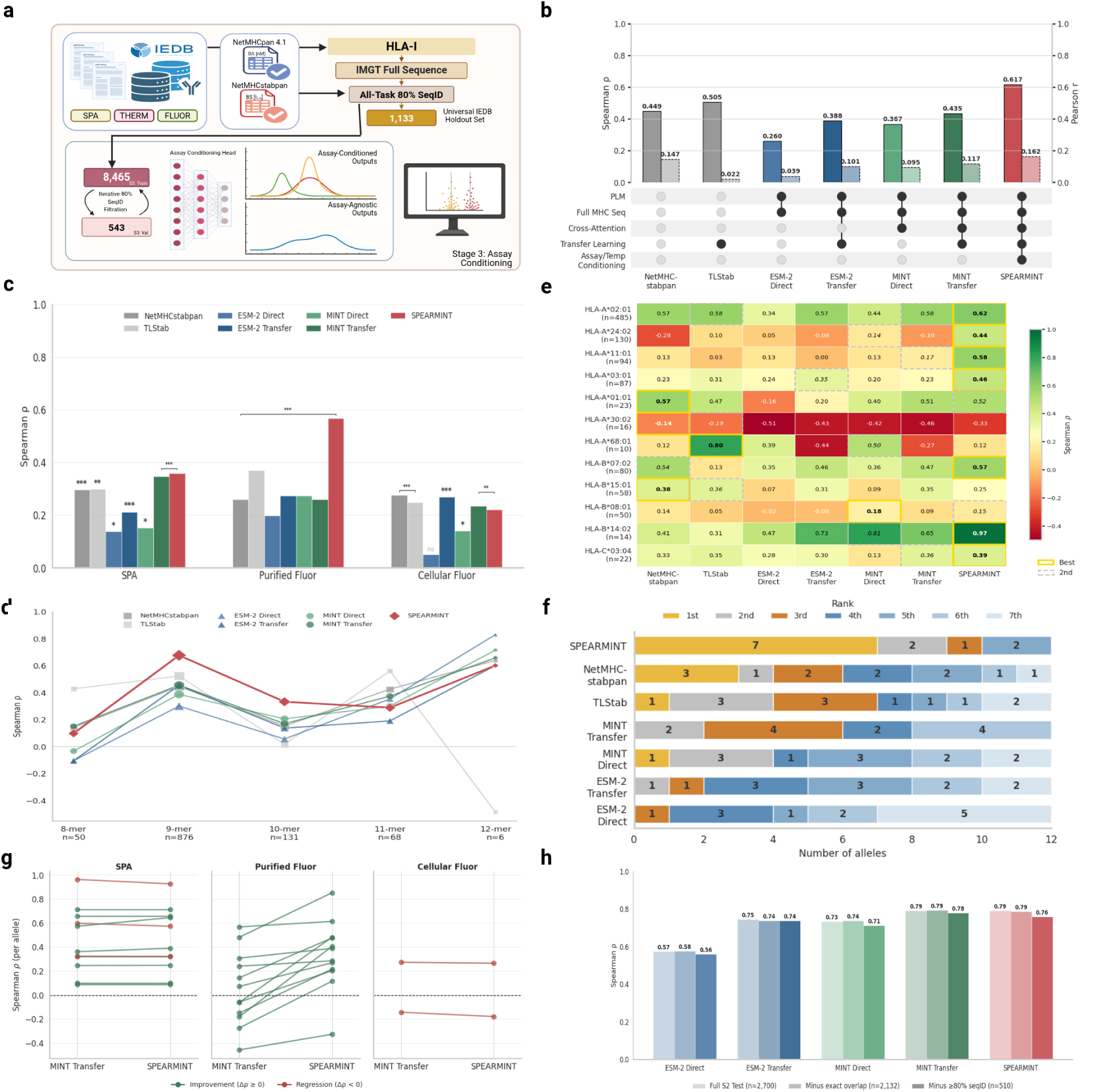
Assay-conditioning mitigates systematic assay shifts and improves pMHC stability prediction across heterogeneous assay contexts. **a.** pMHCs filtered from the previous IEDB heterogeneous assay generalization dataset are recycled for stage 3 training, allowing leakage-controlled re-use of the IEDB holdout data. **b.** Dataset-level benchmark on IEDB holdout with model capabilities annotated. **c.** Assay-stratified benchmark showing Spearman *ρ* for each model. Significance levels are bootstrapped by computing the test statistic (Spearman *ρ*) on permuted labels. **d.** Spearman *ρ* values shown for common k-mers, stratified by model. **e.** Heatmap of per allele Spearman *ρ* values calculated on the IEDB dataset. The top ranking model and runner-up are boxed in the gold and silver boxes per row, respectively. **f.** Distributions of rankings across all alleles for each model. Model order is based on cumulative weighted rank. **g.** Paired dot plots for each assay show allele-level changes in performance between the MINT Transfer model and the flagship SPEARMINT (MINT S3 FiLM) model. Alleles with at least 5 peptides per assay are shown. **h.** Bar plots of models *ρ* values for leakage-controlled partition of the Stage 2 test set, to assess prediction drift during Stage 3 training.

On the universal IEDB holdout, SPEARMINT produced the largest gains on purified fluorescence measurements, reaching *ρ* = 0.567 (*n* = 758), more than doubling MINT Transfer (*ρ* = 0.259) and outperforming TLStab (*ρ* = 0.369) and ESM-2 Transfer (*ρ* = 0.274) as well (**Figure 4b; Supplementary Figure 6a-b**). SPA performance was largely preserved (*ρ* = 0.359 vs. 0.346 for MINT Transfer; *n* = 217), whereas cellular fluorescence remained modest across models (*ρ* = 0.220), consistent with limited training data and noisier cell-based readouts (**Figure 4c; Supplementary Figure 6c**). Across peptide lengths, SPEARMINT achieved the highest Spearman correlation for both 9-mers (*ρ* = 0.675, *n* = 888) and 10-mers (*ρ* = 0.332, *n* = 131), and ranked among the top three models for 10 of 12 qualifying alleles, placing first for 7 (**Figure 4d-f**). In paired allele-level comparisons, assay conditioning improved all 12 evaluated alleles for purified fluorescence (median Δ*ρ* = +0.296; Wilcoxon *p* = 4.9 × 10^−4^), while leaving SPA performance largely unchanged (median Δ*ρ* = 0.000; **Figure 4g**).

To determine whether these gains reflected structured assay adaptation rather than increased model flexibility, we next examined the contributions of temperature, unmodeled inter-protocol variance, and peptide–MHC feature coverage. Unsurprisingly, temperature conditioning improved performance, indicating that incubation conditions contribute measurable signal beyond assay identity alone (**Supplementary Figure 6d**). Within SPA measurements, performance differences between epitopes where the study was seen during training suggest that protocol-specific information is learned implicitly even within SPA data, potentially explaining the universal drop in performance compared to the NetMHCstabpan dataset (**Supplementary Figure 6e**). Finally, stratification by observed anchor residue combinations (P2, PΩ) showed that part of Stage 3 performance is explained by an increased representation of anchor residues per allele, while SPEARMINT nevertheless outperformed MINT Transfer at similar data availability (**Supplementary Figure 6f**). Finally, to ensure that these gains did not come at the cost of corrupting the SPA-trained representation, we evaluated SPEARMINT on the Stage 2 test set after leakage correction and observed only a minor drop in Spearman correlation (0.79 → 0.76), indicating no catastrophic forgetting of the original SPA mapping (**Figure 4h**).

### 3.4 Downstream Transfer to Eluted Ligand and Immunogenicity Prediction

One key application of binding stability prediction is improving downstream immunogenicity prediction. Immunogenicity itself is not a singular phenomenon, but emerges from a cascade of biological filters: MHC binding (binding affinity), stable surface presentation (binding stability), and T-cell recognition, where each step is necessary but not sufficient for the next. Having trained binding affinity and stability predictors in sequence, we evaluated whether stability provides complementary signal to affinity for predicting two downstream endpoints: eluted ligand status (persistent surface presentation) and immunogenicity (T-cell recognition) (**Figure 5a; Supplementary Figure 7a-c**).

**Figure 5:**
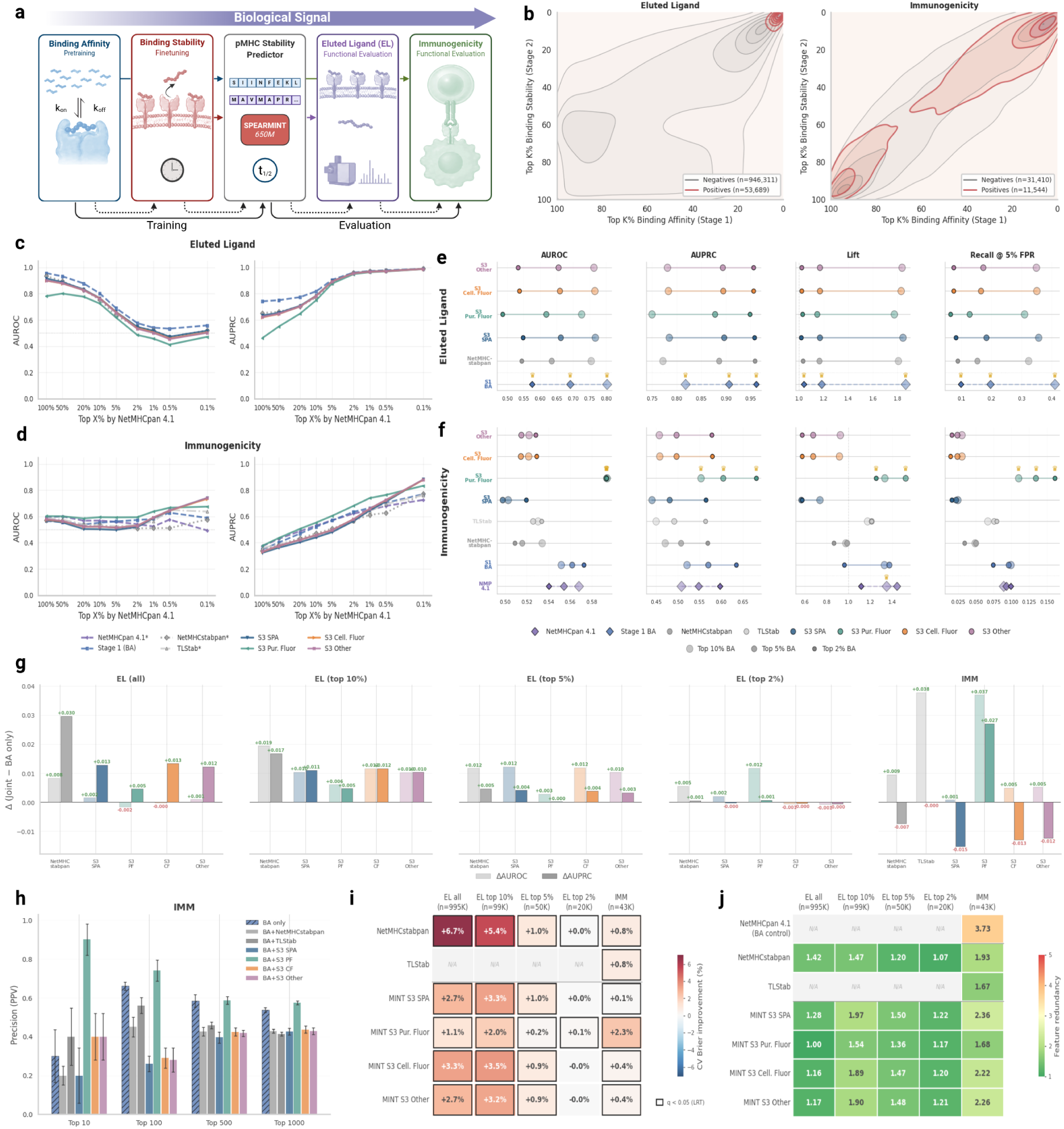
Downstream Functional Analysis. **a.** A biophysical-to-functional signal gradient enables upstream training on pMHC binding affinity and stability, followed by downstream evaluation on eluted ligand presentation and T-cell immunogenicity. **b.** Gaussian KDE plots showing the distributions of positive and negative data points along the percentile rank axes of binding affinity (Stage 1) and binding stability (Stage 2) for EL and IMM datasets. **c-d.** Performance sweeps across NetMHCpan 4.1 predicted affinity thresholds for AUROC and AUPRC. **e-f.** Stratified dot plot demonstrating per-model performance metrics across the predicted top-10, 5, and 2% for EL and IMM dataset partitions. **g.** ΔAUROC and ΔAUPRC between Stage 1 binding affinity model and the joint model trained using each binding stability model shown per stability model. **h.** Precision @K vales shown for immunogenicity dataset per joint model. **i.** Cross-validated Brier score (%) for logistic regression models combining MINT binding affinity with each stability predictor, relative to binding affinity alone (5-fold). Positive values indicate improved calibration. Black outlines denote statistical significance (likelihood ratio test, BH-corrected q < 0.05). **j.** Variance inflation factor (VIF) between MINT binding affinity and each stability predictor, measuring feature redundancy. VIF = 1 indicates no correlation; values above 5 suggest collinearity. NetMHCpan 4.1 BA is included as a correlated control to Stage 1 affinity prediction.

We first examined the distribution of binding affinity and binding stability predictions for each endpoint and found that while the EL positives were tightly clustered about the highest percentile rank of stable binders, the positives for immunogenicity were significantly more disperse (**Figure 5b**). Stratification of peptides by NetMHCpan 4.1 predicted affinities revealed binding affinity to be more predictive of EL outcome over binding stability models at all thresholds of binding affinity (**Figure 5c; Supplementary Figure 7d**). NetMHCpan 4.1 was kept out of the EL evaluation along with TLStab due to unremovable data leakage. However, this trend flipped in the immunogenicity dataset where the assay-conditioned stability models outperformed the baseline stability models as well as affinity prediction models (**Figure 5d; Supplementary Figure 7e**). Of interest, we observed that at the highest-affinity regimes, the more functionally relevant stability assays outperformed the more biophysically controlled ones with cellular fluorescence outperforming purified fluorescence and both outperforming SPA.

Consolidating a handful of affinity gates, and expanding our set of metrics we investigated if there were models or trends that persist across gates and metrics. For EL prediction, the SPA and cellular fluorescence heads performed comparably to NetMHCstabpan (AUROC 0.77 and 0.75, respectively, at the top 10% affinity gate), although all stability models trailed binding affinity on this endpoint (**Figure 5e**). For immunogenicity, however, the purified fluorescence head emerged as the strongest model, achieving AUROC 0.59 and AUPRC 0.60-0.68 across affinity gates, consistently outperforming binding affinity (AUROC 0.55-0.57), NetMHCstabpan (AUROC 0.51-0.53), and TLStab (AUROC 0.53). At the most stringent top 2% affinity gate (n=860, positivity=0.58), purified fluorescence achieved a recall of 0.16 at 5% FPR, more than doubling NetMHCstabpan (0.03) and TLStab (0.08), indicating that assay-aware stability modeling captures immunogenicity-relevant signal not recovered by either affinity predictors or general-purpose stability models (**Figure 5f**).

Finally, to assess whether stability predictions provide additional signal beyond binding affinity, we trained joint logistic regression models combining our Stage 1 binding affinity predictions with each stability predictor via 5-fold cross-validation and measured the change relative to binding affinity prediction alone. For eluted ligand prediction, all stability models yielded consistent gains that peak at intermediate gating levels for AUROC (Δ up to +0.019 for NetMHCstabpan at the top 10% BA gate) and in the ungated setting for AUPRC (Δ up to +0.030), with both metrics converging toward zero at the top 2% gate as EL-positive prevalence approaches one. For immunogenicity, Purified Fluorescence achieved the largest gains (ΔAUROC = +0.037, ΔAUPRC = +0.027), while TLStab matched on AUROC (Δ = +0.038) but not AUPRC, and NetMHCstabpan provided only marginal improvement (ΔAUROC = +0.009, ΔAUPRC = −0.007) (**Figure 5g**). These trends also held when using NetMHCpan 4.1 as the binding affinity predictor, though to a lesser degree (**Supplementary Figure 7f-g**). Top-*K* precision analysis further highlights Purified Fluorescence, which achieves a Precision@10 of 0.90 (ΔP@10 = +0.60 over BA alone) and is the only stability predictor to consistently match or exceed BA-only precision across all cutoffs (**Figure 5h**). In addition, we use likelihood ratio tests, Brier scores [35], and Spearman-based VIF analysis to confirm that stability features provide complementary rather than redundant signal for both endpoints (**Figure 5i-j**).

### 3.5 Retrospective Personalized Neoantigen Prioritization by Antigen Presentation Signal

Having established that assay-conditioned stability predictions improve population-level immunogenicity discrimination beyond binding affinity alone, we next asked whether these gains translate to the clinically relevant setting of personalized neoantigen prioritization, where the goal is to rank candidate neoantigens within individual patients and the candidates are heavily enriched for binding affinity. For this section, we identified four datasets having publicly available datasets with antigen-directed therapies to personalized targets and accompanying immunogenicity data resolved to the epitope level. Across these four independent patient cohorts, we conduct a retrospective personalized neoantigen prioritization analysis spanning multiple treatment modalities, analyzing which presentation signal better predicts experimentally validated T-cell reactivity (**Figure 6a**). Of relevance, synthetic long peptide vaccines, mRNA vaccines, and adoptive T-cell therapies each interface with different parts of the antigen presentation pathway or TCR repertoire, offering unique angles to interrogate the models in the context of the relevant biology (**Figure 6b**). We first characterized each cohort by the number of patients, candidate neoantigens per patient, immunogenicity prevalence, and cohort-level differences in the allele distributions which could directly affect the interpretability of our per-patient metrics (**Figure 6c-e**).

**Figure 6:**
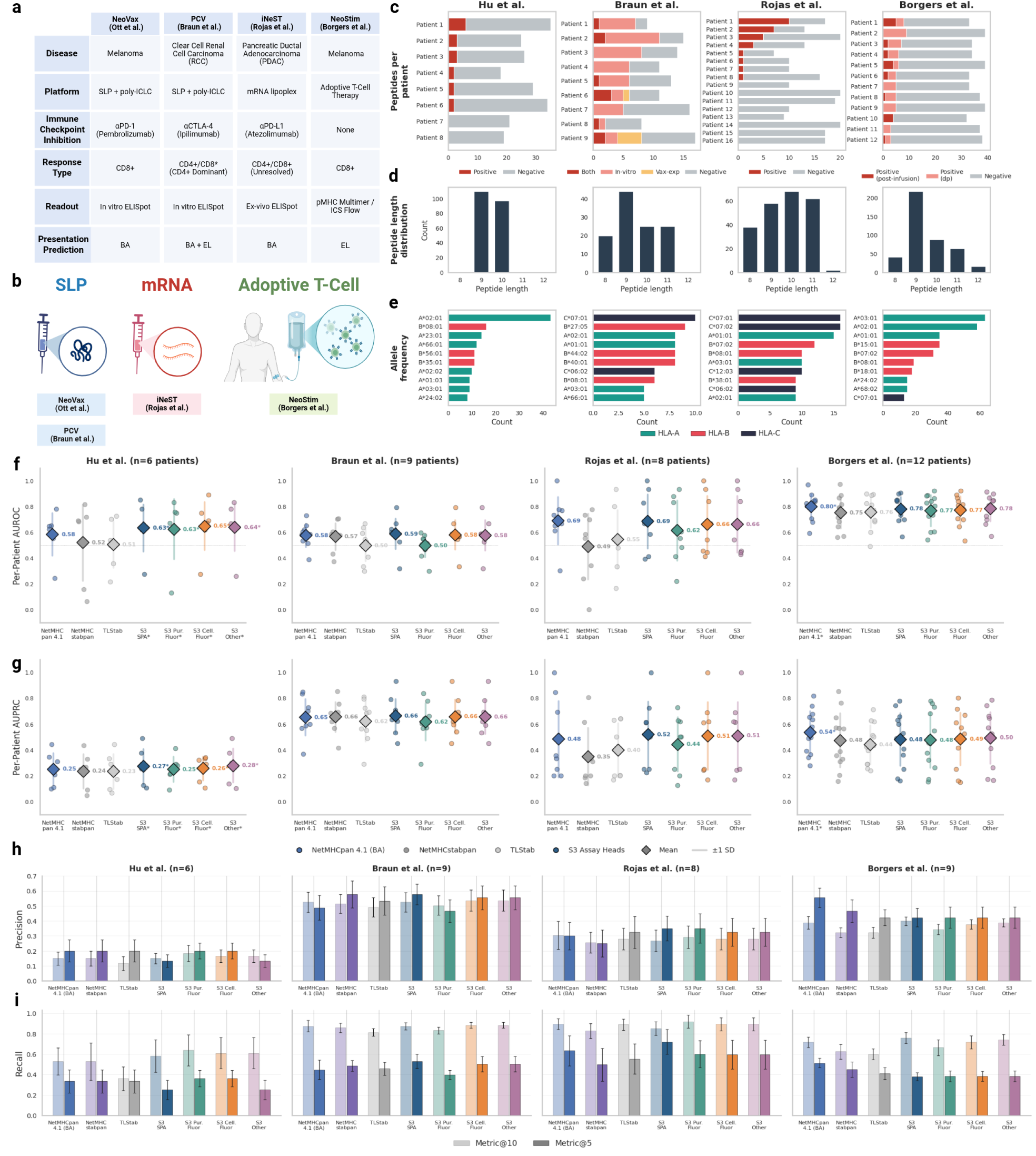
Personalized NeoAntigen Discovery and Validation. **a.** Datasets used in this analysis are presented, characterized by therapeutic, mechanistic, and bioinformatic features. **b.** Visual summary of datasets per therapeutic modality. **c.** Breakdown of peptides tested per patient for each dataset, colored by label (positive=immunogenic, negative=non-immunogenic). **d.** Peptide length distribution per dataset shown aggregated across patients. **e.** Allele frequencies collected across patients shown per dataset. **f-g.** AUROC and AUPRC values are shown per patient for each model along with the medians per model and the interquartile ranges (IQR). **h-i.** Per-patient precision (top) and recall (bottom) at K = 5 (dark) and K = 10 (light) for each stability predictor across four neoantigen cohorts. Metrics are computed per patient and averaged; error bars show standard error of the mean.

Across the four cohorts, SPEARMINT’s assay-conditioned heads match or exceed all baseline models for per-patient neoantigen discrimination, leading on all but one dataset where it came second to NetMHCpan 4.1 (BA) (**Figure 6f-g**). In Hu et al., the Cellular Fluorescence head achieves the highest mean per-patient AUROC (0.65 vs 0.52 for NetMHCstabpan and 0.51 for TLStab) and AUPRC (0.28 vs 0.24 and 0.23, respectively). SPA leads in Braun et al. (AUROC 0.59, AUPRC 0.66) and Rojas et al. (AUROC 0.69, AUPRC 0.52), while in Borgers et al. NetMHCpan 4.1 binding affinity is the strongest single predictor (AUROC 0.80, AUPRC 0.54), with the Other head close behind (AUROC 0.78, AUPRC 0.50) and both outperforming NetMHCstabpan (AUROC 0.75, AUPRC 0.48) and TLStab (AUROC 0.75, AUPRC 0.44). Across all cohorts, SPEARMINT models consistently outperform the general-purpose stability baselines, though no single assay head dominates uniformly, consistent with the heterogeneous experimental conditions underlying each study. These trends held when looking at the precision and recall values at top-*K* of 5 and 10 (**Figure 6h-i**). Of relevance, we found that when NetMHCpan 4.1 incorrectly ranks a positive-negative pair, SPEARMINT heads rescue the correct ordering more frequently than either NetMHCstabpan or TLStab, indicating that SPEARMINT captures complementary discriminative signal rather than mirroring binding affinity (**Supplementary Figure 8a-f**). Interestingly, when restricting evaluation to the more stringent post-administration immunogenicity labels for Braun et al. and Borgers et al., the relative advantage of stability predictions over binding affinity decreases, however more studies are required to explore this pattern further (**Supplementary Figure 8d**).

## 4 Discussion

In this study, we revisit pMHC-I binding stability prediction in light of renewed interest in stability as an underappreciated driver of antigen presentation, immunogenicity [3], and immunodominance [5]. Compared with the mature landscape of binding affinity prediction, public pMHC-I stability prediction remains comparatively narrow, consisting primarily of NetMHCstabpan [9] and TLStab [10], with both sharing similar architectures and the latter exploring novel training strategies to combat substantial data constraints. Due in part to extreme data sparsity, prior benchmarks do not fully resolve the effects of sequence similarity, dataset overlap, and assay-dependent label shifts. As a result, the field has lacked a clear leakage-corrected estimate of quantitative performance on truly unseen peptide–MHC complexes, a gap that becomes especially relevant for non-canonical antigens such as neoepitopes with limited homology to existing training examples. Here, we introduce, to our knowledge, the first protein language model for pMHC-I binding stability prediction and show the benefit of an assay-conditioned prediction approach on a rigorous intra- and inter-task leakage-controlled benchmark. We further demonstrate that predicted half-life carries downstream signal for presentation and immunogenicity beyond binding affinity.

By enforcing similarity-aware splits of our training and test sets across sequential tasks, we aimed to distinguish genuine stability prediction from near-duplicate recognition. Extending the transfer learning approach introduced in TLStab, we present a three-stage curriculum learning approach for peptide:MHC-I (pMHC-I) binding stability prediction with supervised pre-training of binding affinity prediction, fine-tuning on binding stability, and conditioning on assay-specific signals such as assay type and temperature. Using MINT [11], an ESM-2 [12] based protein language model designed with a multimer cross-attention to model the interaction between proteins, and the foundational NetMHCstabpan dataset, we conducted systematic knockouts of the cross attention and the transfer learning, justifying the inclusion of both learning strategy and architectural complexity in robust, leakage-controlled benchmarks.

A second central finding is that the kinetic dissociation of peptide from MHC (half-life) is not an assay-invariant measure of binding stability. For pMHCs measured across multiple stability assays, half-life values showed low rank agreement between protocols, indicating that assay modality introduces shifts in the observed label, as shown in the IEDB examples. In doing so, we observed that all methods exhibited substantial performance degradation, with even the best models falling below a Spearman correlation of 0.4. Our assay-conditioned models address this by treating half-life as a shared latent stability signal observed through protocol-dependent transformations. The pareto optimal performance across all assay measurements, together with preserved SPA performance, suggest that assay conditioning can correct systematic measurement shifts without erasing the original stability representation. More broadly, this result argues for the modeling of immunological measurements as conditional, context-driven observations, rather than direct, assay-independent ground truths.

We next asked if specific assay-conditioned values better correlated with downstream functional immunological endpoints, specifically persistent surface presentation (eluted ligand positivity) and immunogenicity. On the EL task, we show that binding affinity outperformed all stability prediction models across all metrics, while a model trained on both demonstrated the complementary nature of including both measurements. For immunogenicity, however, we demonstrate strong performance of the assay-conditioned models, with cellular fluorescence outperforming purified fluorescence and both outperforming SPA at the most stringent binding affinity gates, reflecting the prior that different experimental measurements of stability may correspond to different functional views of pMHC persistence. These results reconcile the apparent order-dependence of the biology where binding affinity acts as a gate on antigen candidacy and is thus especially predictive of surface presentation, whereas stability ranks persistence within the affinity-filtered set and captures sustained T-cell stimulation potential.

Finally, we test our hypotheses on the stringent task of personalized neoantigen prioritization, a notable outstanding task in the field. Unlike population-level EL or immunogenicity benchmarks, clinical neoantigen cohorts consist of patient-specific candidate sets that have often already undergone extensive computational filtering. The remaining task is therefore not to distinguish obvious binders from obvious non-binders, but to rank a small set of plausible candidates within an individual patient. This makes per-patient neoantigen prioritization a demanding setting for assessing the marginal value of stability. Across four clinical cohorts spanning distinct therapeutic modalities, assay-conditioned stability predictions provided positive per-patient ranking signal and complemented binding affinity, whereas existing general-purpose stability predictors often contributed weak or inconsistent signal. These results suggest that stability can improve neoantigen prioritization, but also emphasize that the relevant stability signal may depend on the therapeutic context, including whether antigens are delivered as synthetic long peptides or encoded by mRNA, and the mechanism by which they are presented.

Several limitations of this study remain. First, unlike many recent pMHC modeling pipelines, we did not use EL labels during pre-training. This was intentional: EL is a downstream composite readout, integrating processing, binding, stability, abundance, and MS detectability. While EL supervision may improve representation quality for antigen presentation, it can also blur the distinction between kinetic stability and downstream presentation likelihood. We therefore opted to treat EL as a functional evaluation endpoint. Another limitation is that assay-conditioned stability data remain sparse and imbalanced, particularly for cellular fluorescence measurements, limiting the extent to which cell-based stability behavior can be learned and generalized. In the same vein, even though we observed a boost in performance for unseen pMHCs of studies that appeared in training, we do not model additional variables beyond assay type and temperature. This is a conscious choice, owing to the curse of dimensionality, though we suspect this gap in performance can be meaningfully reduced with additional well-curated data. Third, our models are sequence-based and do not encode any structure information or auxiliary signals for antigen presentation including abundance or proteasomal processing and transport. Lastly, a key limitation is that although our framework allows for the handling of HLA allele mutants, due to only a single mutant for a minority of related peptides being present in the available data, it was excluded from analysis. Taken together, these limitations point to the next advances in stability prediction coming from not only architectural and training modifications but also richer and diverse data across different stratifying metadata as well.

Future progress rests on treating pMHC stability not as a correlate of binding affinity, but rather an integral component of a broader antigen-presentation model that integrates binding, processing, abundance, and surface persistence. Paired multi-assay measurements of the same peptide–MHC complexes would help disentangle intrinsic stability from assay-specific measurement effects. Increased collection of data across diverse HLA alleles would increase performance on less-represented alleles, critical for translational utility. Similarly, on the modeling side, measures of uncertainty quantification would have immediate benefit by allowing researchers to act on a particular predicted stability in accordance with its associated confidence score. Additionally, given the prevalence and significance of CD4+ responses in the neoantigen setting, extending these models to MHC-II data would be a high-yield next step. Overall, by combining leakage-aware benchmarking, supervised transfer learning, assay-conditioned recalibration, and downstream functional evaluation, we establish pMHC half-life as a distinct biophysical signal connecting peptide–MHC binding to antigen presentation and T-cell recognition. This work provides a foundation for next-generation antigen-presentation models that are not only more accurate, but more measurement-aware and functionally grounded, establishing assay-conditioning as a step toward more calibrated binding stability prediction and reliable personalized neoantigen prioritization.

## Data and Code Availability

All sequence data (peptide:MHC pairs), computed results, and code used for training and evaluating SPEARMINT can be found on: https://github.com/pirl-unc/spearmint. For ease of use, the model and tokenizer for SPEARMINT can be found on Hug-gingFace at: https://huggingface.co/dkarthikeyan1/spearmint. Additionally, the Stage 1 binding affinity models and Stage 2 MINT Transfer models may be found at https://huggingface.co/dkarthikeyan1/mint-stage1-affinity and https://huggingface.co/dkarthikeyan1/mint-stage2-stability respectively.

## Acknowledgments and Disclosure of Funding

This work would not have been possible without numerous fruitful conversations. We thank Michael Bryan, Elian Belot, Matt Dean, and the team at Serova for early conversations on the direction of this work. We thank Dr. Colin Raffel for his generous sharing of compute resources. We thank Andy Lee, Sara Peterson, and our lab at PIRL for helping to review our manuscript. Finally, we are grateful to all the stewards of the public data commons for their generous and valuable contributions, whose effort make our study and countless others possible.

This work was supported largely by the National Science Foundation Graduate Research Fellowship. The authors have no competing interests to declare.

## Supplementary Figures

**Supplementary Figure 1:**
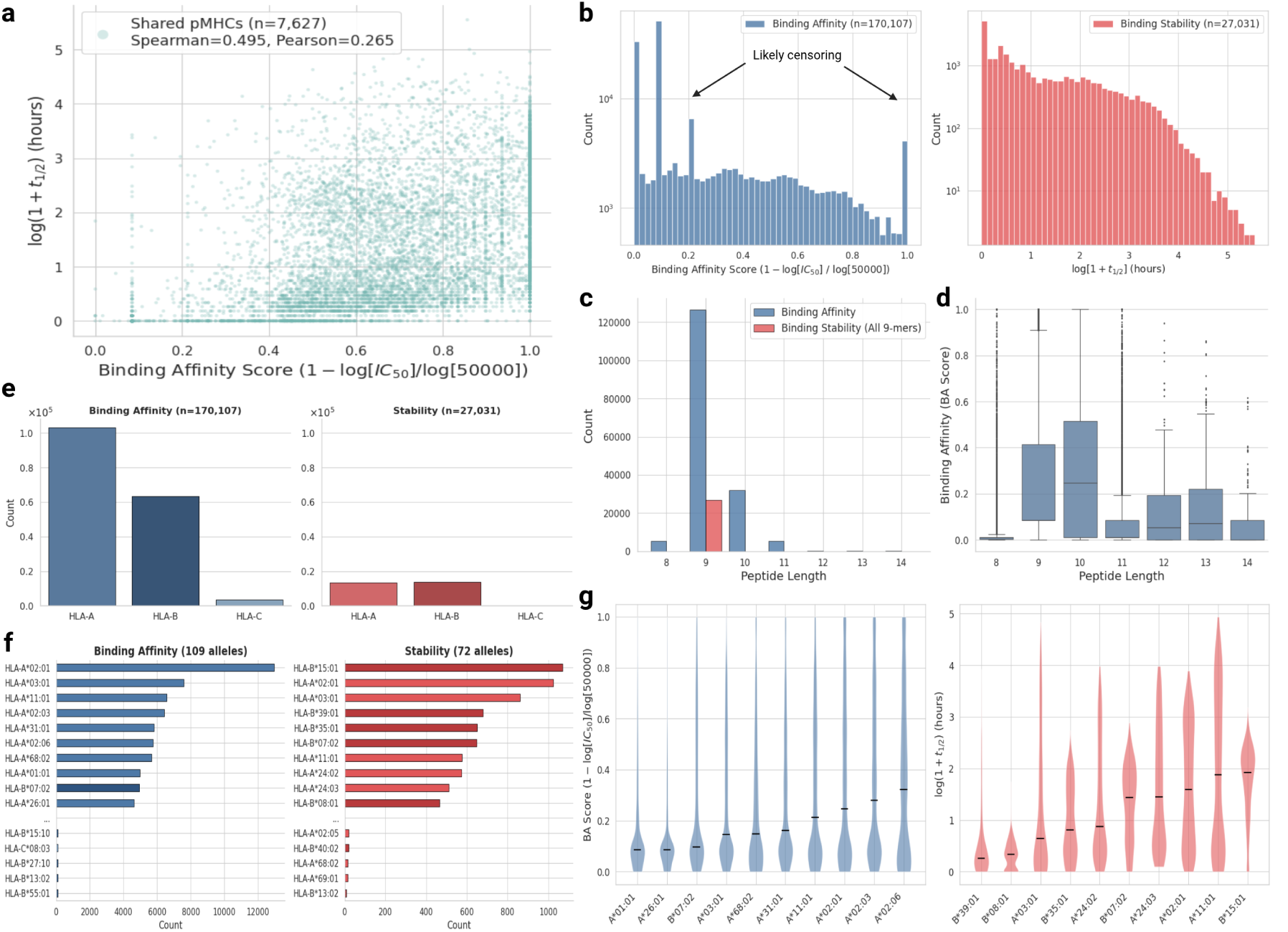
Dataset composition of binding affinity and binding stability data. **a.** Binding affinity / stability correlation between shared pMHCs in NetMHC4.1 (BA) and NetMHCstabpan data (n=7627). **b.** Label distribution of binding affinity (blue) and binding stability data (red). Likely censoring events are highlighted for binding affinity data. **c.** Bar plot of peptide length distributions for binding affinity and binding stability (9-mer only). **d.** Violin plot showing binding affinity values by peptide length. **e.** HLA distribution of binding affinity and binding stability data (Note: binding stability has no HLA-C data). **f.** Per-allele breakdown of binding affinity and binding stability data (top-10 and bottom-5 are shown). **g.** Allele-specific distributions of binding affinity and binding stability, sorted in increasing order among the top-10 most represented alleles for each dataset.

**Supplementary Figure 2:**
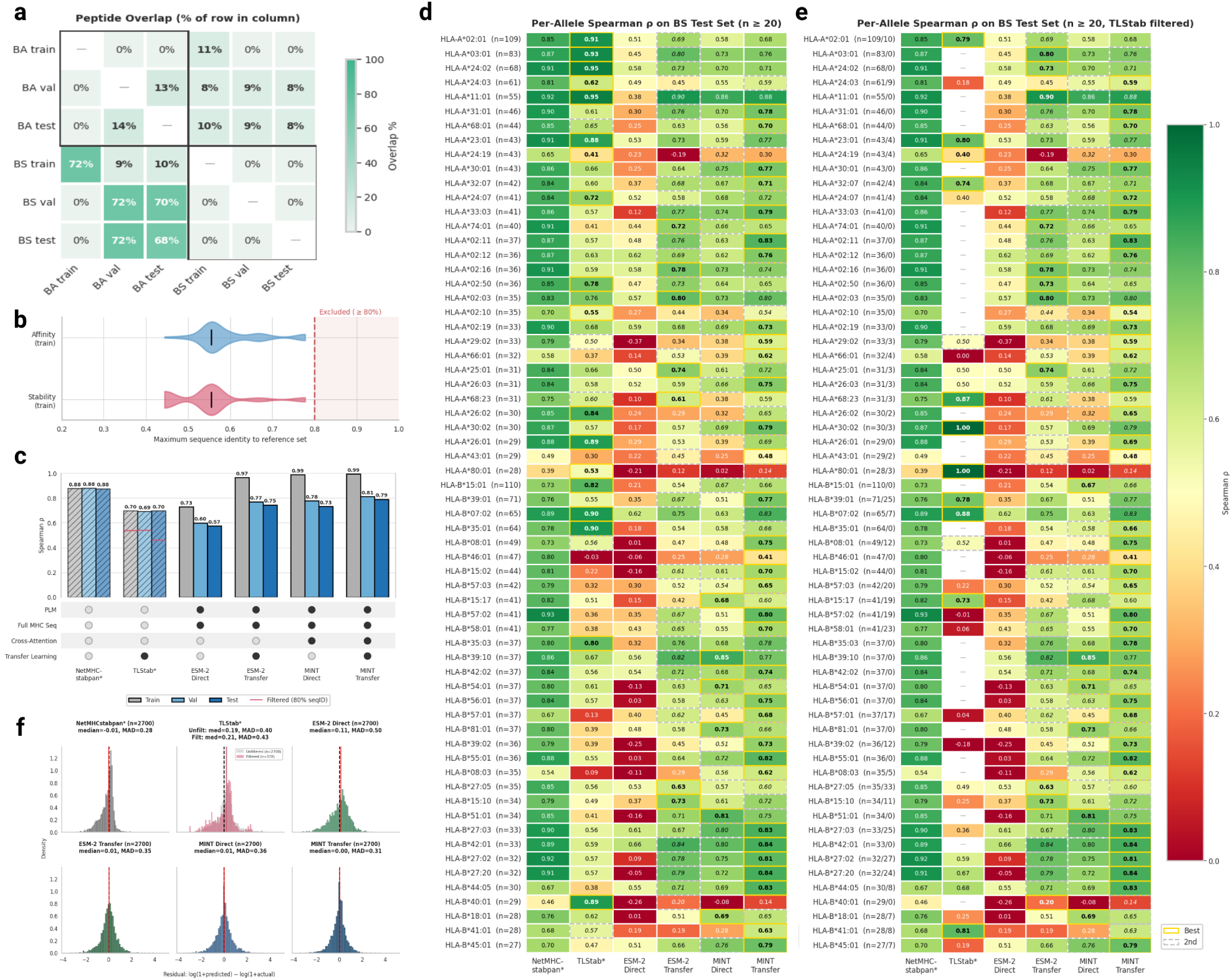
Robustness analysis for NetMHCstabpan dataset results. **a.** Intra-and Inter-task peptide overlap between the train/test/val splits in the affinity and stability tasks. **b.** Sequence identity of the holdout ‘test’ set to pMHCs in the ‘train’ sets for both binding affinity (blue) and binding stability (red). **c.** Per-split Spearman *ρ* of all models, annotated by complexity of architecture and training. **d-e.** Heatmap of per-allele Spearman *ρ* values for the unfiltered and TLStab-filtered holdout sets. NetMHCstabpan-excluded first and second-place rankings are reported under each model. **f.** Distribution of residuals per model with annotated means and mean absolute deviations (MAD). Filtered (red) and unfiltered (gray) holdout sets shown for TLStab.

**Supplementary Figure 3:**
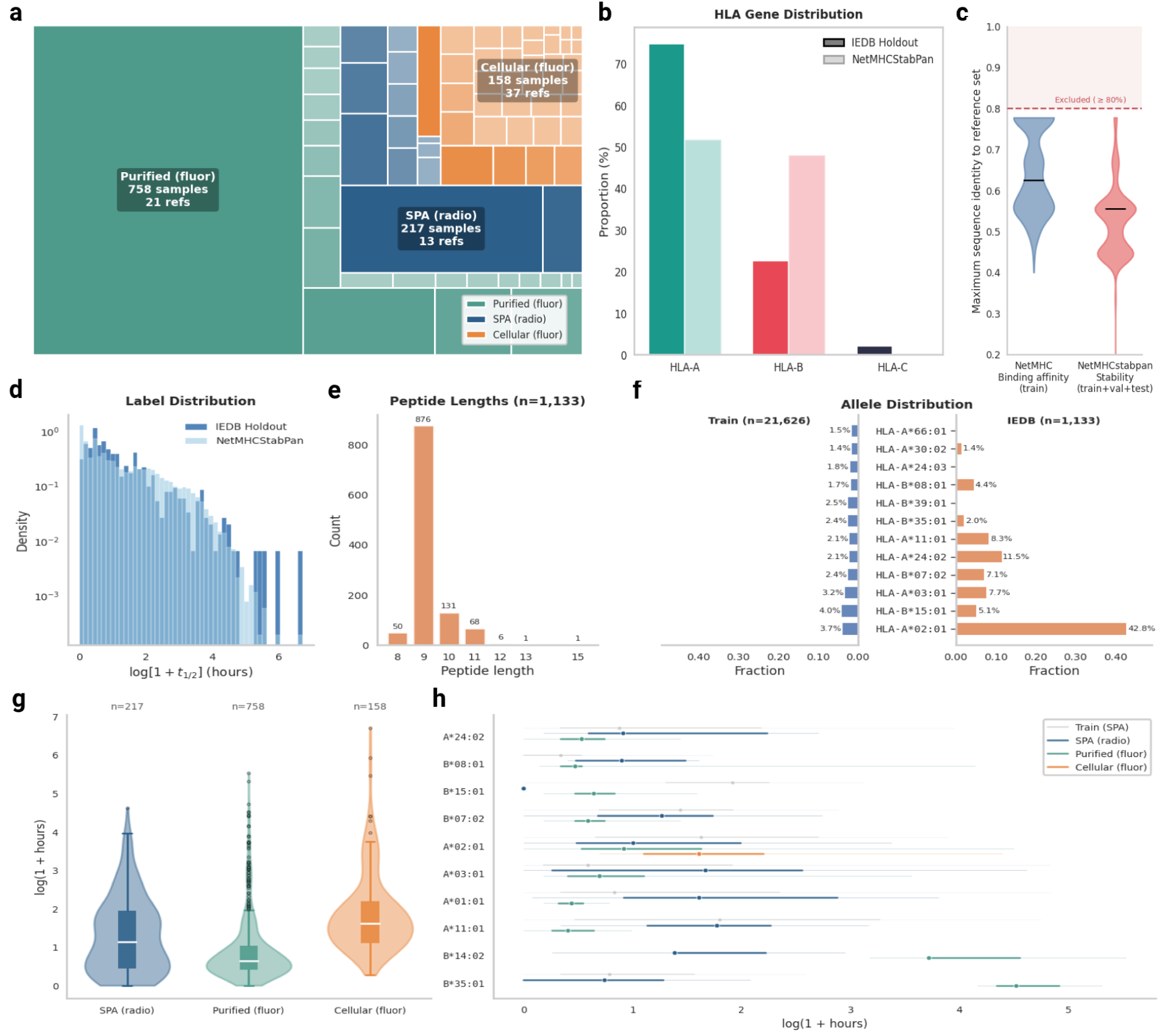
Dataset characterization of heterogeneous IEDB dataset. **a.** Intra- and inter-task peptide overlap between the train/test/val splits in the affinity and stability tasks. **b.** HLA gene-level distribution analysis between NetMHCstabpan training data and the IEDB holdout test set. **c.** Sequence identity of the holdout ‘test’ set to pMHCs in the ‘train’ sets for both binding affinity (blue) and binding stability (red). **d.** Distribution of labels between NetMHCstabpan and IEDB holdout test set. **e.** Bar chart of peptide length counts for IEDB held out data. **f.** Allele distributions of NetMHCstabpan and IEDB holdout. **g.** IEDB holdout label distributions, stratified by determining assay. **h.** Assay-level distributions further stratified by presenting HLA allele for the top-10 most represented alleles.

**Supplementary Figure 4:**
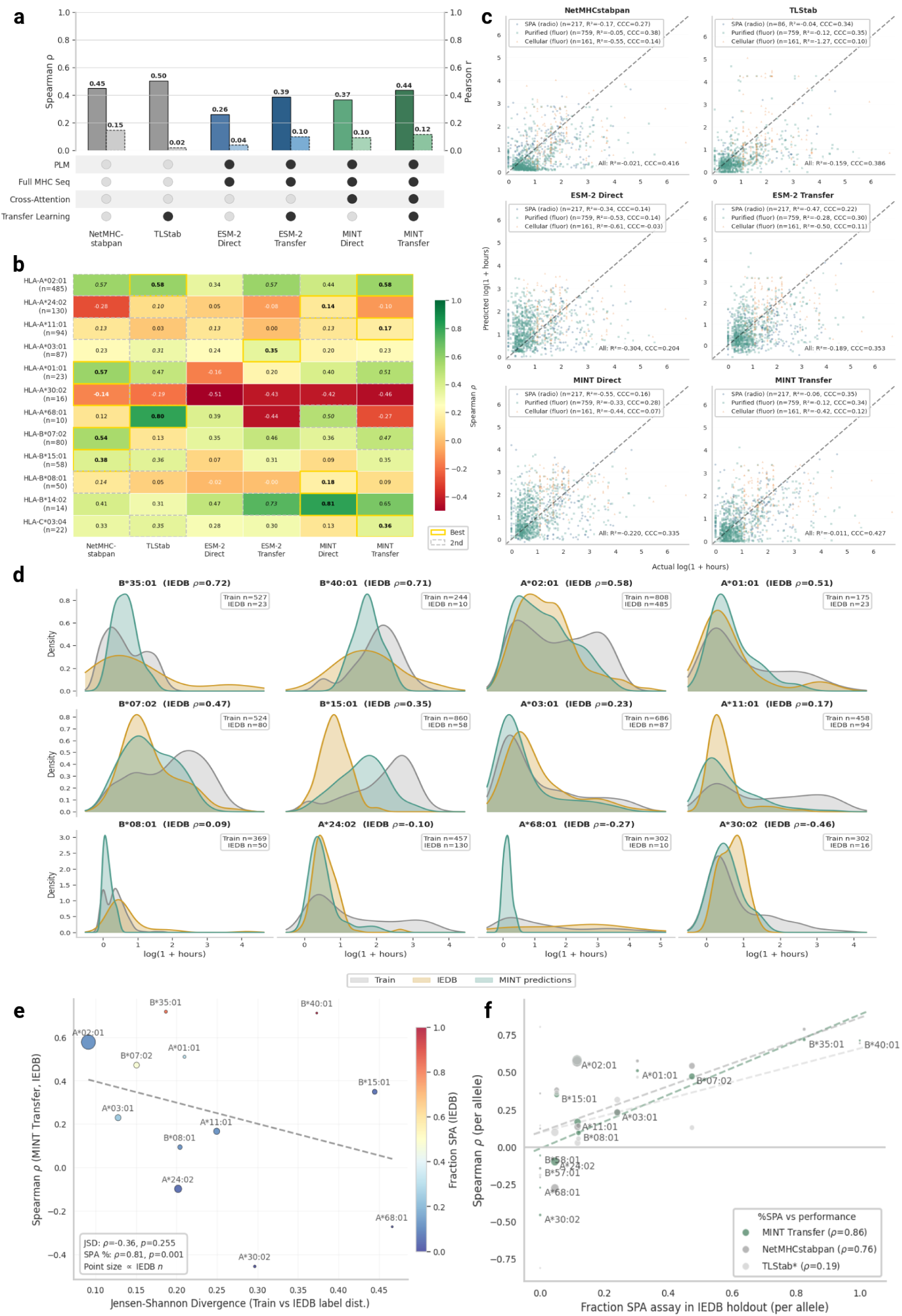
Confounding variable analysis for heterogeneous IEDB dataset. **a.** Upset plot of model performance across entire IEDB test set. **b.** Heatmap of per-allele Spearman *ρ* values for each model. **c.** Scatterplot of predicted and actual log half-life values stratified by assay-type. **d.** Distribution shift plot showing true labels, predicted values, and training set values for top-12 most represented alleles. **e.** Correlation plot between model performance and JS divergence in training vs. evaluation labels, plotted per-allele for MINT Transfer. **f.** Correlation plot between model performance and proportion of examples per-allele coming from the NetMHCstabpan SPA dataset shown per model.

**Supplementary Figure 5:**
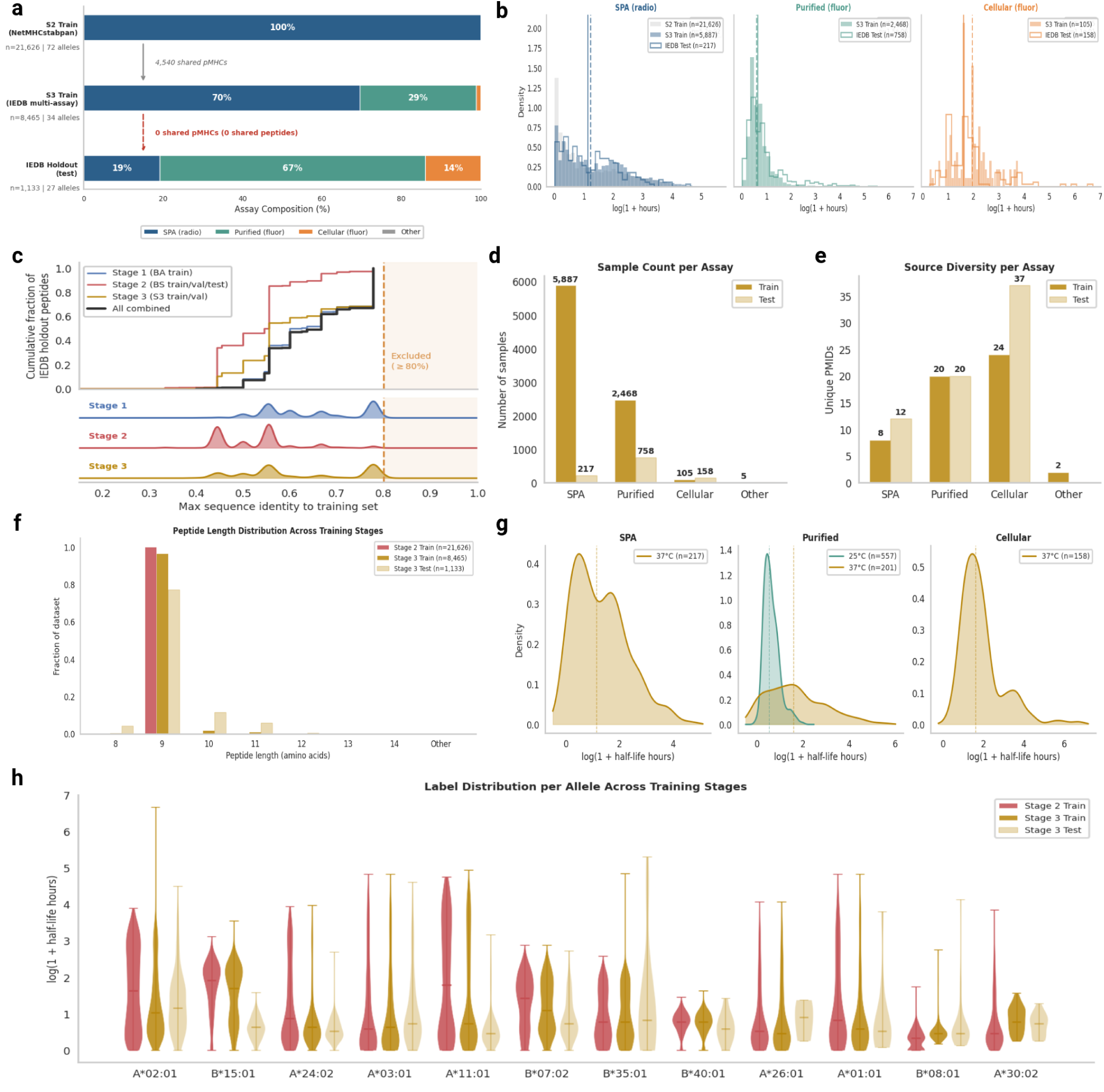
Dataset composition of IEDB assay-conditioning data. **a.** Frequency bar plot showing the originating assay and overlap between examples for Stage 2 Train, Stage 3 Train, and the universal IEDB holdout test set. **b.** Per-assay label distributions for the different assay types within each IEDB split (train vs test). **c.** Sequence similarity leakage plot showing maximum sequence identity per IEDB test peptide and every upstream data source (stage 1, stage 2, and stage 3). **d.** Sample count per assay type in IEDB. **e.** Unique studies per assay type in IEDB. **f.** Peptide length distribution across training stages. **g.** Density plot of labels shown per-assay, stratified by temperature. **h.** Allele-specific label distributions across training stages.

**Supplementary Figure 6:**
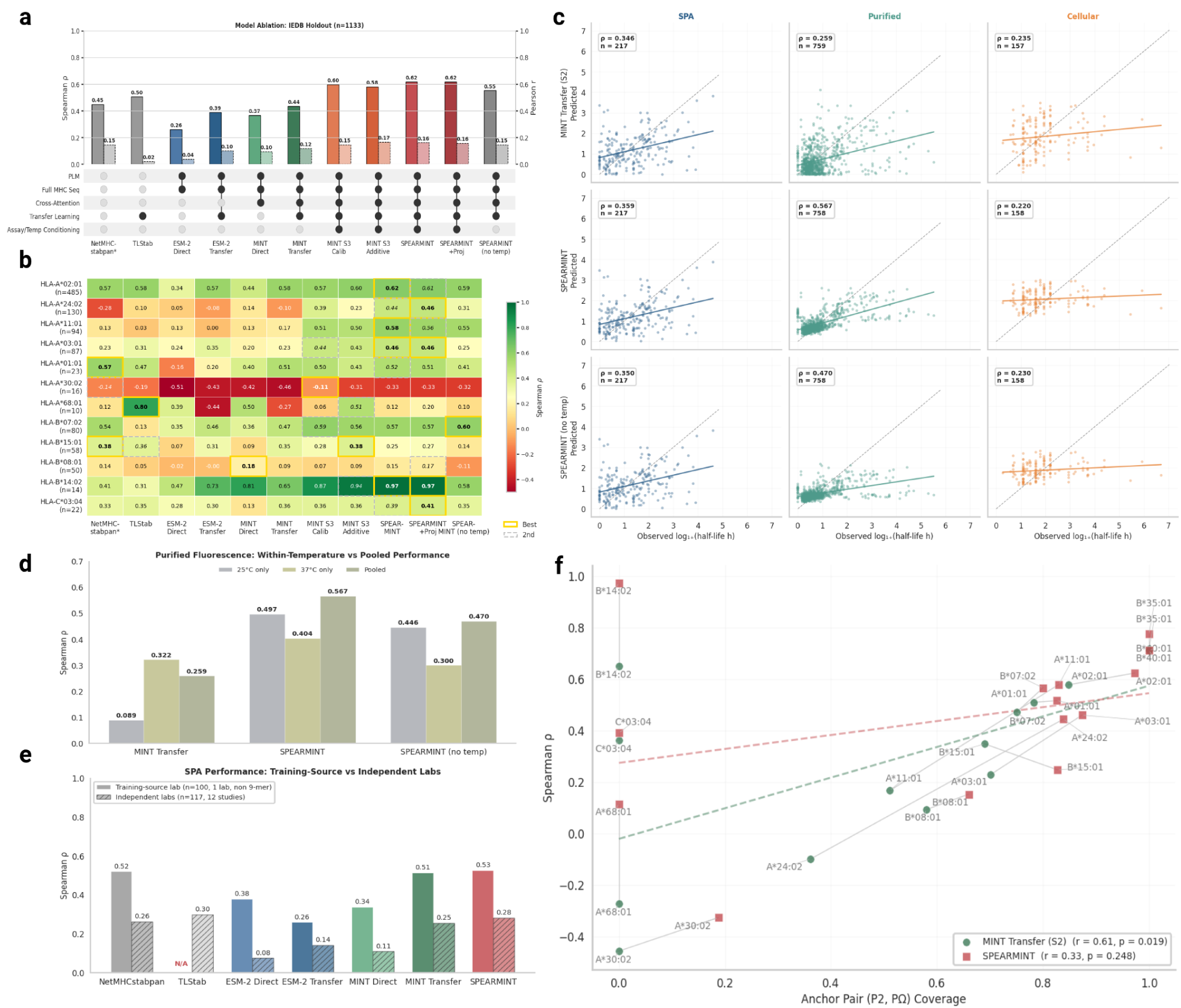
Assay-conditioning improves per-assay and overall performance on a mixed-assay dataset. **a.** Full model performance plot showing baseline models as well as all three assay conditioning variants and two ablation variants on the most performant conditioning method (FiLM [32]). **b.** Per-allele heatmap of Spearman *ρ* values for all baseline models and assay-conditioning variants. **c.** Scatterplot of predicted versus observed half life values of IEDB test peptides split by assay x model. **d.** Barplots showing temperature-conditioned performance (Purified Fluorescence assay-only). **e.** Seen lab vs. unseen lab generalization benchmark. TLStab is ‘N/A’ due to near complete overlap of data with its training set. **f.** Unseen anchor residue pair generalization benchmark. Per-allele spearman performances plotted against anchor position coverage (fraction of test pairs seen in training).

**Supplementary Figure 7:**
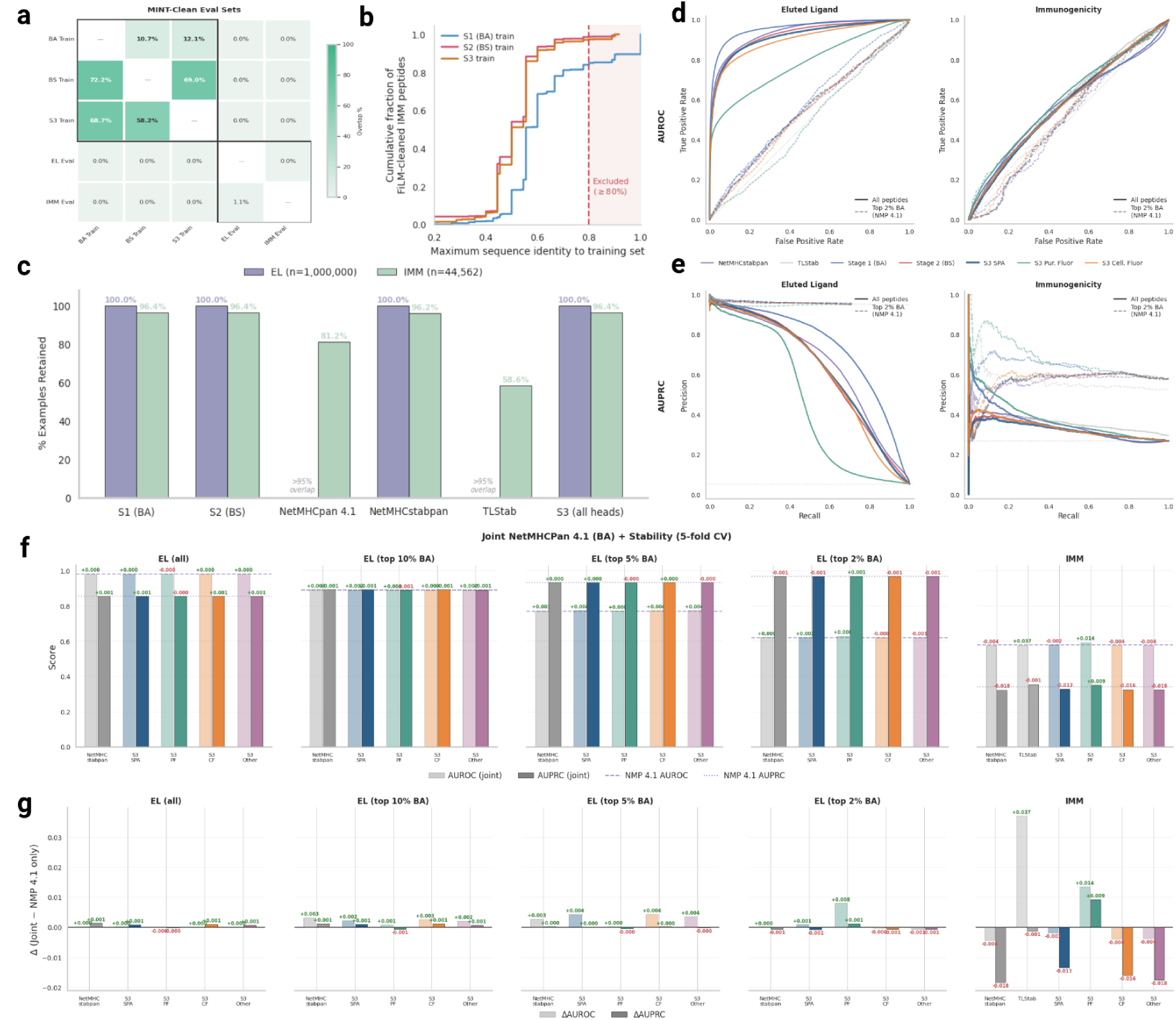
Extended downstream analyses. **a.** Heatmap showing the percent overlap between the three-stage training and downstream evaluation datasets (% of row in column). **b.** ECDF of sequence identities of each evaluation set to the three training stage corpora. **c**. Barplots showing the number of examples retained after model specific leakage filtering. Only exact peptide overlaps are removed. **d-e.** Four panel plot showing the AUROC and AUPRC curves for EL and IMM datasets. Curves for the top 2% of binders (as determined by NetMHCpan 4.1) are shown with dashed lines. **f.** Bar plots showing the AUROC and AUPRC of joint binding affinity model and stability prediction models (logistic regression) with the NetMHCpan 4.1 (BA) model. **g.** ΔAUROC and ΔAUPRC barplots between the joint model and NetMHCpan 4.1 alone.

**Supplementary Figure 8:**
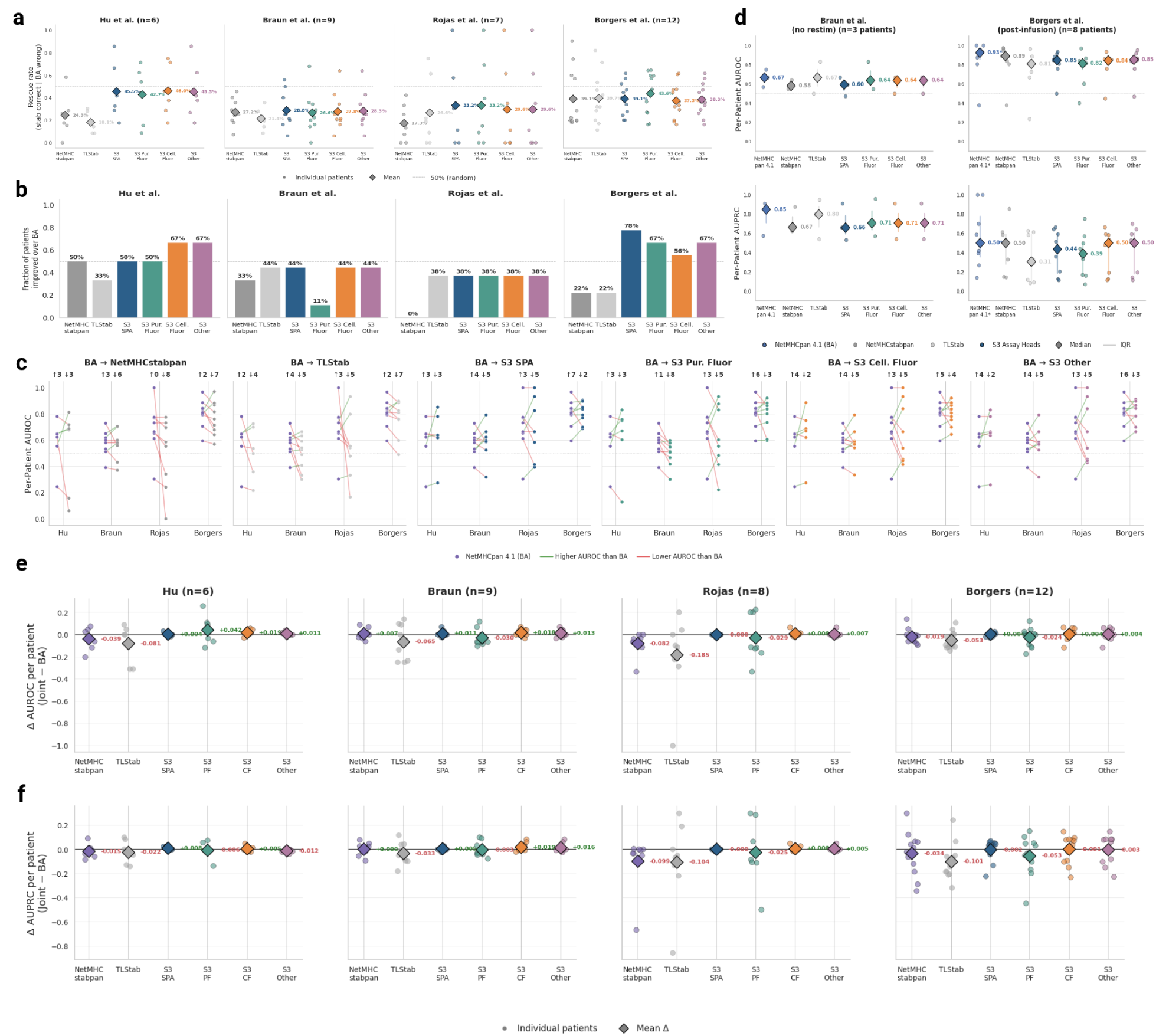
Robustness analysis for personalized neoantigen dataset results. **a.** Fraction of NetMHC4.1-misranked peptide pairs correctly re-ranked by each stability model, per patient. **b.** Summary plot showing the fraction of patients per dataset whose AUROC performance increased over NetMHCpan 4.1 for each stability model. **c.** Paired dotplot showing AUROC performance changes between NetMHCpan 4.1 and all stability models for each dataset. **d.** AUROC and AUPRC values shown per patient for Braun and Borgers on the no-stim and long lived response labels, respectively. **e-f.** ΔAUROC and ΔAUPRC shown per patient for the joint (BS + BA) and NetMHCpan 4.1 (BA) model. Models were fit using logistic regression on predicted values on the IMM dataset and transferred to the per-patient neoantigen setting.

