## Supplementary Information for "Peptide:MHC Binding Stability Prediction Using Protein Language Models"

### SPEARMINT

**D Checkpoint Selection** **11**

Correspondence to {dkarthikeyan1, alex.rubinsteyn}@unc.edu.

### A Data Curation

#### A.1 Binding Affinity Data

Binding affinity (BA) training data were obtained from the NetMHCpan 4.1 [1] training corpus, comprising 208,093 peptide:MHC-I measurements across five data tranches. Labels are continuous scores derived from measured $\mathrm{IC}_{50}$ values via the standard log-transformation [2]:

$$\mathrm{BA}_{\text{score}} = 1 - \frac{\log(\mathrm{IC}_{50})}{\log(50{,}000)} \tag{1}$$

yielding values in $[0, 1]$, where higher scores indicate stronger binding.

Allele names were normalized to two-field resolution (e.g., HLA-A*02:01) and mapped to full-length MHC-I $\alpha$-chain amino acid sequences ($\sim$365 residues) obtained from the IPD-IMGT/HLA database [3]. Non-human alleles (H-2, BoLA) and alleles lacking full-length consensus sequences were discarded, reducing the dataset from 208,093 to 170,107 measurements across 109 HLA-I alleles. An initial 80/10/10 random split was followed by a strict peptide-level deduplication step: 968 peptide sequences appearing in both the training and validation/test partitions were identified and all corresponding training rows (9,402) were removed and redistributed equally to the validation and test sets. This yielded a final BA dataset of 126,683 training, 21,712 validation, and 21,712 test examples, with zero peptide overlap between training and evaluation partitions.

#### A.2 Binding Stability Data

Binding stability (BS) measurements were obtained from the NetMHCstabpan training corpus [4], comprising 28,166 peptide::MHC-I complexes with half-life labels ($t_{1/2}$) measured in hours. Of particular interest, the dataset consists exclusively of 9-mer peptides bound to their HLA class I molecules. While the data was generated from multiple studies, they were all conducted by the same lab, using the same method: scintillation proximity assay (SPA) [5]. The same allele normalization and full-length MHC sequence mapping procedure was applied, yielding 27,034 measurements across 72 HLA-I alleles. Stability labels span a wide dynamic range ($t_{1/2} \in [0, 256.7]$ hours) with a large number weak binders concentrated around $t_{1/2} < 1$. To stabilize training, we used a log-transform:

$$\mathrm{BS}_{\text{score}} = \log(1 + t_{1/2}) \tag{2}$$

We chose this transformation over the fitted stability score $s = 2^{-t_0/t_{1/2}}$ used by NetMHCstabpan [4], due to the latter's requirement of a threshold parameter $t_0$ and propensity to saturate near $s = 1$ for long-lived complexes, compressing the dynamic range of the label space. In comparison, the log-transform is parameter-free, preserves discrimination across the full dynamic range, and yields an unbounded target compatible with the linear output of our regression head. Predictions are mapped back to native half-life units via $\hat{t}_{1/2} = \exp(\hat{y}) - 1$ at inference time.

#### A.3 Independent IEDB Heterogeneous Holdout

To evaluate generalization beyond the NetMHCstabpan data distribution, we constructed an independent holdout set of pMHCs and their measured stability half-lives from the Immune Epitope Database (IEDB) [6]. We downloaded all (positive and negative) MHC class-I binding stability measurements from IEDB (10,896 records as of March 2026) and retained only valid peptides composed of the 20 standard amino acids. All peptides were within 8-15 amino acids in length. To this subset, we normalized allele names to the standard two-field format, including resolution of serological designations (e.g., HLA-A2 $\rightarrow$ HLA-A*02:01) which affected less than 1% of the data, strictly limited to HLA-A1, HLA-A2, HLA-A3, HLA-A11, HLA-B8, and HLA-B27, which were all given the canonical ':01' denominations except for HLA-B27 which was resolved as HLA-B*27:05. Once we had the canonical two field nomenclature, we map these to full-length MHC sequences and apply mutations at the designated positions. Finally, to preserve the label distribution, we converted half-life measurements to hours instead of the default minutes, while accounting for a few direct submissions that were uploaded in hours and noted in the assay comments [7, 8]. To prevent data leakage from the binding affinity and stability datasets, we removed all peptides sharing $\geq$80% sequence identity

(Levenshtein distance) with any peptide in the combined binding affinity and stability training corpora, reducing the evaluation set to 1,187 measurements. Where multiple measurements existed for the same peptide–allele pair (57 of 1,130 unique pairs), we retained the median half-life, yielding a final holdout set of 1,130 unique pMHC complexes across 27 HLA-I alleles for the assay-agnostic approach.

##### A.4 Assay-Conditioned Training Data

To train the assay-conditioned Stage 3 model heads, we returned to the full IEDB binding stability export (10,896 records). After applying quality filters (MHC class I restriction, valid peptides of 8–15 standard amino acids, allele normalization and MHC sequence mapping, and retention of quantitative half-life measurements), 10,391 records remained. We binned each measurement into one of four assay groups based on the IEDB “Assay - Method” field: *SPA* (radioactivity-based, purified MHC), *Purified Fluorescence* (fluorescence-based, purified MHC), Cellular Fluorescence (fluorescence-based, cell-surface MHC), and *Other*. Incubation temperatures were extracted from the IEDB assay comments field where available; when absent, we assigned defaults of 37°C for SPA, Purified Fluorescence, and Cellular Fluorescence assays, and 25°C for iTopia direct submissions [7, 8]. Half-life units were converted to hours as described previously.

Where multiple measurements existed for the same (peptide, allele, assay, temperature) combination, we retained the median half-life. We then performed peptide-level decontamination, removing any remaining peptide with $\geq$80% sequence identity (normalized Levenshtein distance) to any test-set peptide, to prevent near-duplicate leakage. The cleaned pool was split into training and validation partitions at an 85/15 ratio, stratified by assay type to ensure representation of each protocol in both splits. A final iterative decontamination pass identified validation peptides with $\geq$80% identity to any training peptide and migrated them to the training set, repeating until convergence. Zero (peptide, MHC sequence) overlap exists between any pair of splits.

The resulting Stage 3 dataset comprises 8,465 training entries (5,357 unique peptides, 34 alleles), 543 validation entries (512 unique peptides, 15 alleles), and 1,133 held-out test entries (813 unique peptides, 27 alleles). The training set is dominated by SPA measurements (5,887; 69.5%), followed by Purified Fluorescence (2,468; 29.1%) and Cellular Fluorescence (105; 1.2%), reflecting the underlying distribution of stability assays in IEDB. Temperature coverage is 74% at 37°C and 26% at 25°C.

##### A.5 Downstream Functional Evaluation

To evaluate the impact of stability prediction as an auxiliary signal to binding affinity for peptide prioritization in real-world settings, we benchmarked all models on two common and mechanistically relevant functional readouts: eluted ligand (EL) identification and T-cell immunogenicity (IMM). Of importance, neither signal was used at any stage of training for our models. For both tasks, models were evaluated in a zero-shot transfer setting: predicted stability scores were used directly as ranking features with no additional fine-tuning or threshold calibration on the evaluation data. This design tests whether pMHC stability, as a biophysical intermediate between binding and antigen presentation, carries signal that transfers to downstream functional outcomes without task-specific supervision.

###### A.5.1 Eluted Ligand Data

Our eluted ligand (EL) dataset was taken from the NetMHCpan 4.1 training corpus [1]. The raw data comprise 13M pMHCs across five cross-validation folds , where labels indicate whether a peptide was identified as an MHC-I ligand by mass spectrometry (1) or is a decoy (0) drawn from the proteome. We first filtered to human HLA alleles, discarding all non-human MHC data, reducing the dataset to 3,679,405 records across 130 HLA-I alleles. Allele names were normalized to two-field resolution and similarly mapped to full-length MHC-I sequences as described throughout. To make evaluation tractable while preserving the class distribution, we performed a stratified random subsample to 1,000,000 peptide–allele pairs, maintaining the original positive rate of 5.4% (53,689 positives, 946,311 negatives).

##### A.5.2 Immunogenicity Data

Our immunogenicity benchmark datset was drawn from the IEDB T-cell assay database [6] (export dated March 2026; 122,913 records) to which the following filters were applied: (1) only $CD8^+$ T cell responses were kept, (2) alleles with at least the serotype resolution (e.g., HLA-A2) were retained (3) peptides of 8–15 amino acids in length and (4) only those composed of the 20 standard amino acids were kept. Records were then aggregated by unique (peptide, allele) pairs, yielding $n_{\text{positive}}$ and $n_{\text{negative}}$ response counts per pair. A binary label was assigned as positive if $n_{\text{positive}} > 0$. Allele normalization and MHC sequence mapping (as described above) removed 3,878 entries (8.0%) with unmappable alleles, yielding a final dataset of 44,562 unique peptide–allele pairs.

##### A.5.3 Personalized Neoantigen Therapeutics Data

We finally posit personalized neoantigen prioritization as not only a translationally relevant problem, but also one of the most stringent of evaluation conditions, fundamentally distinct from population or dataset-level metrics. Not only is the evaluation done on a per patient basis, these candidate peptides have also already undergone extensive computational filtering such that the remaining discrimination task is confined to an already-enriched pool from which conventional filters have removed the easy negatives, providing a trenchant assessment of the marginal contributions of stability prediction.

Thus, to assess whether stability predictions generalize to the clinically relevant task of neoantigen prioritization, we curated a panel of four published neoantigen directed clinical cohorts spanning distinct cancer types, assay modalities, and patient populations. Cohorts were selected according to three criteria: (1) availability of patient-level HLA typing at two-field resolution or better, (2) experimentally validated binary immunogenicity labels for mutant neoepitopes presented on autologous HLA alleles, and (3) sufficient per-patient sample size (multiple pMHCs per patient, at least one immunogenic and non-immunogenic peptide per patient, and at least five patients) to permit per-patient AUROC computation.

Our search yielded four datasets with descriptions drawn from their methods:

- **NeoVax (Ott et al. 2017 & Hu et al. 2021)** [9, 10]: NeoVax is a personalized neoantigen vaccine platform that uses long-peptides formulated with poly-ICLC adjuvant that was administered to high risk melanoma patients subcutaneously (s.c.) on days 1, 4, 8, 15 and 22 (priming phase) and weeks 12 and 20 (booster phase). The Hu et al. study is a follow up of the former and explores 6 + 2 patients' (six previously reported in Ott et al.) long term responses and single-cell phenotype trajectories to the treatment regimen. Immunogenicity of individual peptides are determined using after one round of in vitro stimulation by IFN-$\gamma$ ELISpot using week 16 post-vaccination PBMCs. In this dataset while persistent memory responses were detected *ex vivo* for $CD4^+$ T cells, all CD8 responses were in vitro stimulated, with some being developing into memory responses. The raw data for this dataset was taken from Hu et al.'s Supplemental Dataset 4a and the label used was the `'Peptide pulsed autologous APC (16 weeks)'` column.
- **PCV (Braun et al. 2025)** [11]: This study introduces the personalized cancer vaccine or PCV, which is the same platform at the Ott et al. NeoVax, aimed towards patients with clear cell renal cell carcinoma (RCC), of whom some are taking checkpoint inhibitor. In this study, individualized synthetic long peptides (SLPs) were similarly administered subcutaneously as well as intradermally on days 1, 4, 8, 15 and 22 for vaccine priming and weeks 12 and 20 for the booster phase. In this study the authors report a number of measures of immunlogical reactivity presented in their Supplementary Table 2: `'InVitro_PeptideStim_ReplicateXX'`, `'InVitro_NoStim_ReplicateXX'`, `'VaccineExpanded_TumorStim_ReplicateXX'`, `'VaccineExpanded_NoStim_ReplicateXX'`. Of importance the paper's reported values were taken from the in vitro section of the table. Due to the sparsity of the vaccine-expanded measurements, we use the in vitro immunogenicity readouts for this dataset. Among the available label definitions we adopt the most permissive, calling a peptide immunogenic if its in vitro peptide-stimulated response is significantly elevated over the paired no-stim control by the source's reported two-sided $t$-test (`Ttest_pvalue_InVitroStim` $< 0.05$), without imposing an additional fold-change threshold. Where several overlapping SLPs tile the same mutation and share one predicted minimal epitope, we collapse them to a single entry per (patient, HLA allele,

minimal epitope) by logical OR, treating an epitope as immunogenic if any SLP context elicited a response.

- **iNeST (Rojas et al. 2023 & Sethna et al. 2025)** [12, 13]: In this study, the authors follow a regimen of surgery on resectable tumors, anti-PDL1 given on week six after surgery, their autogene cevumeran (an individualized vaccine based on mRNA-lipoplex nanoparticles encoding up to 20 MHCI and MHCII restricted neoantigens) given as seven weekly priming doses beginning on week 9 (boosters on week 17 and 46), and 12 cycles of mFOLFIRINOX starting on week 21. The source material can be found in Rojas et al's Supplementary Table 5 under the column `'ELISpot Response'`. Of note, their vaccine payload is specifically spleen targeting. Additionally, this study's supplemental notes that while these responses were *ex vivo* responses, their assay based on [14] does not distinguish between CD4+ and CD8+ responses and operates on the best predicted MHC-I allele and best predicted MHC-I epitope per spanning neoantigen sequence. Thus the results for this dataset are based on the predicted epitopes to bonafide positivities of unkown ontology (CD4+ or CD8+).
- **NeoStim (Borgers et al. 2025** [15]: Unlike the other treatments in this group, NeoStim stands by itself as an adoptive cellular therapy and not a vaccine. Instead, the NeoStim drug product uses multimers for a handful of personalized-neoantigens to identify and enrich for antigen-specific T cells that are reintroduced into the patient in higher numbers and a more activated state. Positivity against specific neoantigens was determined using functional readouts of IFN-$\gamma$ and/or TNF and/or CD107a. The source material for this dataset were taken from the study's Supplementary Data Set 1 where the labels included were: `'Detected in Vessel'`, `'Detected in DP'`, `'Detected pre-infusion'`, and `'Detected Post-infusion'`.

Each cohort was harmonized into a common schema containing mutant and wild-type peptide sequences, two-field HLA allele identifiers, and binary immunogenicity labels taken from each study's reported results and supplementary information. For cohorts with multiple label definitions (Braun: stimulated, unstimulated, $\geq$55% threshold; Borgers: drug product response, vessel response), we report results under the label with the highest positivity rates used by the original study (i.e. with re-stimulation). NetMHCpan 4.1 binding affinity, NetMHCstabpan half-life, and TLStab predictions were computed for each mutant peptide to serve as baselines.

### B Model

#### B.1 Tokenization and Input Representation

Both MINT and the ESM-2 baseline share a common tokenization scheme standardized in the MINT paper [16]. Mapping their scheme to our work, peptide and MHC-I sequences are tokenized independently: each sequence is prepended with a <cls> token and appended with an <eos> token, then padded to the maximum sequence length per sequence type for the batch. The two tokenized chains are concatenated along the sequence dimension to form a single input tensor:

$$\mathbf{x} = [\texttt{<cls>},\ p_1, \ldots, p_l,\ \texttt{<eos>},\ \texttt{<pad>}_{\ldots},\ \texttt{<cls>},\ m_1, \ldots, m_p,\ \texttt{<eos>},\ \texttt{<pad>}_{\ldots}]$$

with its corresponding chain_id tensor:

$$\mathbf{c} = [0_1, \ldots, 0_L, 1_1, \ldots, 1_M]$$

where $p_1, \ldots, p_l$ and $m_1, \ldots, m_p$ denote the peptide and MHC residue tokens, of length l and p, respectively padded out to length L and length P. No explicit separator token is inserted between the two chains. A parallel chain-ID tensor assigns identifier 0 to all peptide positions and identifier 1 to all MHC positions (including their respective special tokens and padding).

For MINT, these chain IDs are consumed by the cross-chain attention mechanism: the attention mask is modified so that tokens attend only within their own chain by default, with cross-chain interactions introduced through designated attention heads. For the ESM-2 single chain baseline, chain IDs are ignored and the model processes the concatenated input as a single sequence, with the <eos>/<cls> tokens at the chain boundary serving as the only implicit delimiter. After the final transformer layer, <cls>, <eos>, and <pad> tokens are masked out, and the remaining residue-level representations are mean-pooled across both chains to produce a fixed-dimensional embedding ($d = 1{,}280$), which is passed to the projection head for regression on either binding affinity or binding stability (**Supplementary Figure 1a**).

#### B.2 Architecture

**ESM-2.** ESM-2 [17] is a protein language model pre-trained via masked language modeling on sequences from UniRef50 [18] that has been become widely used as an out-of-the-box sequence featurizer for many downstream tasks, given a number of favorable properties of the model that emerge from large-scale training and sufficient parameters. There are various sizes of the models in terms of their parameter counts ranging from 8M to 15B, with the 650M model (33 transformer layers, hidden dimension $d = 1,280$) being a common choice for its performance and accessibility. Notably, ESM-2 learns contextual residue-level representations that implicitly encode protein structure with attention maps that recover inter-residue contacts without structural supervision, strong performance on capturing evolutionary fitness landscapes [19], as demonstrated by competitive zero-shot prediction of mutational effects from deep mutational scanning experiments, and many other biophysical properties that have been observed using mechanistic interpretability methods [20, 21]. These properties have demonstrated success in the peptide-MHC binding affinity space [22] and have yet to be demonstrated in the binding stability setting. We hypothesize that these properties make ESM-2 a natural backbone for pMHC binding prediction, where both structural complementarity and residue-level interaction may be captured in the attention maps (Supplementary Figure 1b).

**MINT.** MINT [16] modifies the ESM-2 650M architecture to explicitly model protein-protein interactions through a dual-attention mechanism. MINT's pre-training consists of masked language modeling on $\sim$96M PPIs from the STRING database [23], where the model learned to distinguish intra- from inter-chain co-evolutionary signals using chain-aware masking.

Architecturally, each transformer layer in MINT contains two independent multi-head attention modules with separate intra- and inter-chain $\mathbf{Q}$, $\mathbf{K}$, $\mathbf{V}$ projections (Supplementary Figure 1c). Given the chain-identity tensor $\mathbf{c}$ (where $c_i = 0$ for peptide tokens and $c_i = 1$ for MHC tokens), a binary mask $M_{ij} = \mathbb{I}[c_i \neq c_j]$ partitions all token pairs into intra-chain ($M_{ij} = 0$) and cross-chain ($M_{ij} = 1$) subsets. Raw attention logits are assembled by selecting intra-chain logits from the first module and cross-chain logits from the second:

$$\alpha_{ij} = (1 - M_{ij})\, \alpha_{ij}^{\text{intra}} \;+\; M_{ij}\, \alpha_{ij}^{\text{cross}} \tag{3}$$

A single softmax is applied over the assembled logits, and value aggregation routes each position through the corresponding module's value projection:

$$\mathbf{o}_i = \sum_{j:\, c_j = c_i} \tilde{\alpha}_{ij}\, \mathbf{V}_j^{\text{intra}} \;+\; \sum_{j:\, c_j \neq c_i} \tilde{\alpha}_{ij}\, \mathbf{V}_j^{\text{cross}} \tag{4}$$

where $\tilde{\alpha}$ denotes the softmax-normalized attention weights. This dual-attention mechanism doubles the attention parameters per layer but enables the model to learn fundamentally different interaction patterns for intra-molecular context and inter-molecular contacts.

**Downstream head.** For the downstream stability prediction task, we append a two-layer projection head to the MINT/ESM backbone. After the final transformer layer, <cls>, <eos>, and <pad> tokens are masked out, and the remaining residue-level representations are mean-pooled across both chains to produce a fixed-dimensional embedding ($\mathbf{h}_{\text{pool}} \in \mathbb{R}^{1280}$). The projection head computes:

$$\hat{y} = W_2\ \text{Dropout}\big(\text{ReLU}(W_1 \mathbf{h}_{\text{pool}} + b_1)\big) + b_2 \tag{5}$$

where $W_1 \in \mathbb{R}^{d_h \times 1280}$, $W_2 \in \mathbb{R}^{1 \times d_h}$, and $d_h$ is the projection hidden dimension. Of importance, given that the output is unbounded, we do not use a sigmoid output layer, which would have been more appropriate for a model trained on NetMHCstabpan's normalization of stability values to [0,1].

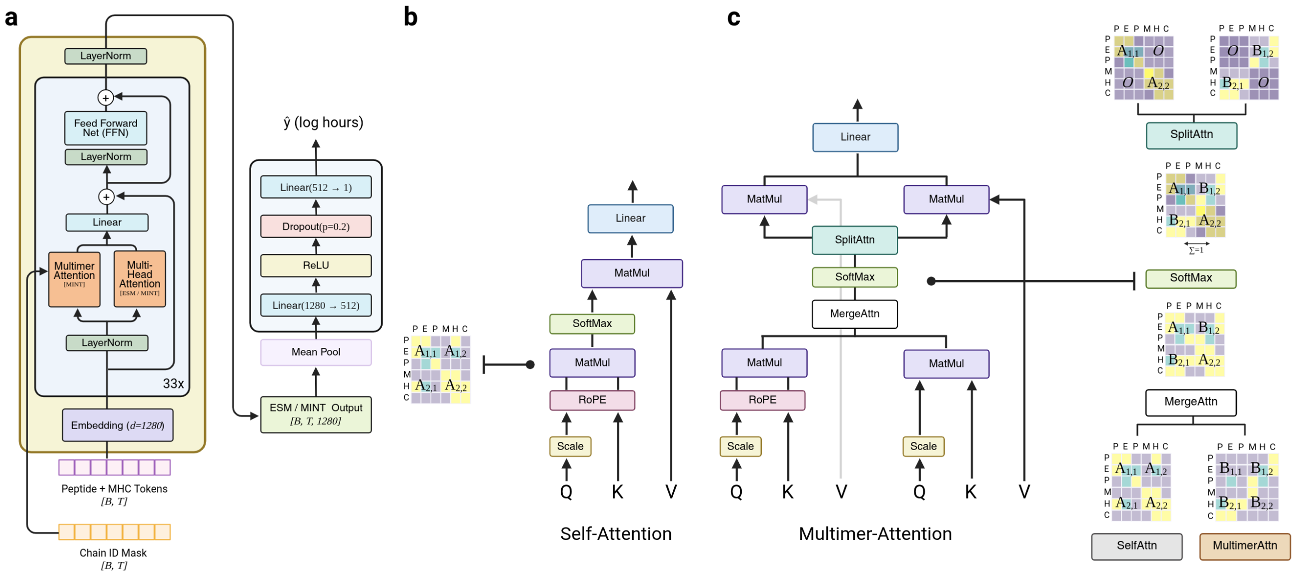

**Supplementary Figure 1:** *Model Architecture.* **a.** Spearmint architecture diagram. Main ESM-2/MINT backbone and projection layer are separated for clarity. Inspired by [24]. **b.** Self-attention as used by the ESM-2 and MINT architectures with corresponding attention map over unified peptide:MHC input. **c.** Multimer-attention diagram showing the independent attention operators and interaction attention maps.

### C Assay Conditioning

The IEDB [6] aggregates data from various labs who deposit their experimental results along with thte relevant and accepted metadata. While invaluable as a resource for increasing the data available to researchers, the accompanying heterogeneity it comes with can prove to be itself a challenge. Stability data in the IEDB aggregates measurements from heterogeneous experimental protocols: scintillation proximity assays (SPA), purified-MHC fluorescence, cellular surface-display assays, among others each measure biophysically distinct cellular phenomena which we observed to introduce systematic shifts in the measured half-lives of the same pMHC. To account for this, we explored the use a lightweight conditioning module on top of the frozen Stage 2 model, designed to transform the learned NetMHCstabpan distribution to distributions derived from the various assay types.

#### C.1 Setup

Our use of a conditioning module, as opposed to retraining the projection layer or additional layers of the MINT backbone were motivated in part by a strong prior on the interoperability of binding stability assays and, more practically, by the fact that there were not many examples to train on. Based in our observation that that the Spearman rank of pMHCs measured using different assay-types was low, indicating that the assay protocol could result in changes to not only absolute, but also relative binding stability, we sought to learn these assay-specific distribution shifts via exploring increasingly complex operators. Specifically, we evaluated three conditioning architectures of increasing expressivity: (i) a per-assay affine calibration with a small residual MLP ( 134K parameters), which assumes assay differences are primarily scale and offset shifts; (ii) a latent residual fusion module ( 559K parameters), which learns a residual correction in the hidden representation space; and the established (iii) Feature-wise Linear Modulation [25] (FiLM; 810K parameters), which produces sample-specific, feature-wise scale and shift vectors conditioned jointly on assay type, temperature, and the pMHC representation itself (Figure 2a-c).

To encode the assay type we assigned every example with a categorical assay label $a \in \mathcal{A} = \{\texttt{SPA}, \texttt{Purified\_Fluor}, \texttt{Cellular\_Fluor}, \texttt{Other}\}$. While there were only two temperatures in our training data and we could have encoded these as low vs high temperature conditions, we decided to stick with a continuous incubation temperature $T$ (°C; defaulting to 37°C when unreported).

These are then consumed by the each of the individual conditioning layers as a learned assay embedding $\mathbf{e}_a = \mathrm{Embed}(a) \in \mathbb{R}^{d_a}$ and a linear temperature projection $\mathbf{e}_T = W_T T + b_T \in \mathbb{R}^{d_T}$. Of note, we ensured that all approaches began from an identity initialization, such that at the start of Stage 3 training, the model exactly reproduces the frozen Stage 2 prediction $\hat{y}^{(S2)}$. This guarantees that training begins from a known-good prediction surface, and the conditioning parameters need only learn a residual correction. In all three modes the MINT backbone is fully frozen; only the conditioning parameters, metadata embeddings, and (optionally) the Stage 2 projection head are trained. We go into the finer grained details of each conditioning method below:

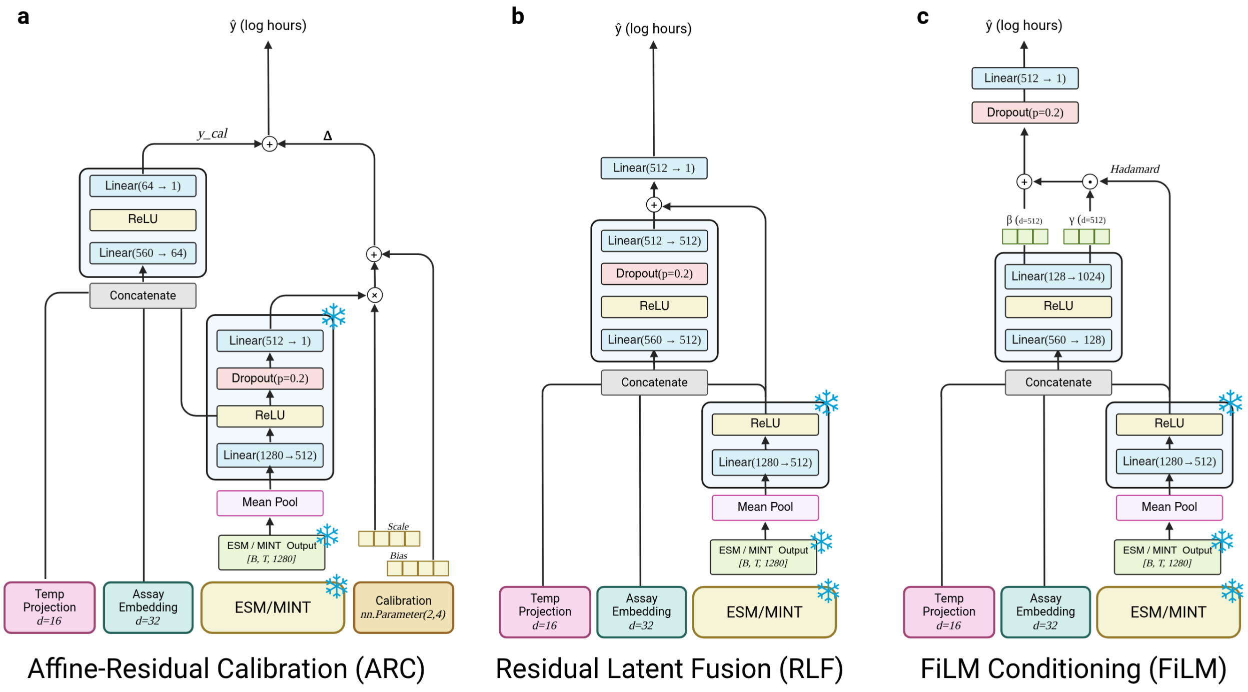

**Supplementary Figure 2:** *Assay Conditioning.* **a.** Spearmint architecture diagram. Main ESM-2/MINT backbone and projection layer are separated for clarity. Inspired by [24]. **b.** Self-attention as used by the ESM-2 and MINT architectures with corresponding attention map over unified peptide:MHC input. **c.** Multimer-attention diagram showing the independent attention operators and interaction attention maps.

#### C.2 Methods

##### C.2.1 Affine-Residual Calibration

The calibration variant operates directly on the frozen Stage 2 scalar output $\hat{y}^{(S2)}$ rather than on the hidden representation. Each assay type $g$ is assigned a learnable scale $\alpha_g$ and bias $\beta_g$, initialized to $\alpha_g = 1$, $\beta_g = 0$. A small residual MLP provides an additional correction term:

$$\hat{y} = \alpha_g\, \hat{y}^{(S2)} + \beta_g + \underbrace{\mathrm{MLP}_{\mathrm{res}}\big([\mathbf{h};\, \mathbf{e}_a;\, \mathbf{e}_T]\big)}_{\Delta} \tag{6}$$

where $\mathrm{MLP}_{\mathrm{res}}$ maps from $\mathbb{R}^{d_h+d_a+d_T}$ through a hidden layer of dimension $d_{\mathrm{res}} = 256$ (ReLU, optional dropout) to a scalar $\Delta$. The output layer is zero-initialized, so $\Delta = 0$ at initialization and the model recovers the Stage 2 prediction exactly.

This is the most parameter-efficient variant at $\sim$134K trainable parameters with default dimensions. The inductive bias is that assay differences are primarily scale and offset shifts, with the residual MLP handling any remaining nonlinear corrections. In this variant, the entire Stage 2 pipeline: backbone, projection head, *and* readout is frozen. Only the per-assay affine parameters, metadata embeddings, and residual MLP are trained.

##### C.2.2 Residual Latent Fusion (Additive)

In the latent residual variant, a residual vector is computed from the concatenation of $\mathbf{h}$, $\mathbf{e}_a$, and $\mathbf{e}_T$ and added directly to the hidden state:

$$\mathbf{r} = \mathrm{MLP}_{\mathrm{fuse}}\big([\mathbf{h};\, \mathbf{e}_a;\, \mathbf{e}_T]\big) \in \mathbb{R}^{d_h}, \qquad \hat{y} = \mathrm{readout}(\mathbf{h} + \mathbf{r}) \tag{7}$$

where $\mathrm{MLP}_{\mathrm{fuse}}$ is $\mathrm{Linear}(d_a + d_T + d_h \to d_h)$, $\mathrm{ReLU}$, optional dropout, and $\mathrm{Linear}(d_h \to d_h)$. The output layer of the fusion MLP is zero-initialized so that $\mathbf{r} = \mathbf{0}$ at the start of training, recovering the Stage 2 prediction. An optional post-fusion dropout is applied before the readout. With the projection frozen and default dimensions ($d_a = 32$, $d_T = 32$), this variant has $\sim$559K trainable parameters and is taken from the principle that residual connections throughout a neural network can trivially learn the identity function and form a function class that supersets the previous layer's learned mapping, which would be useful to retain Stage 2 behavior.

##### C.2.3 FiLM (Feature-wise Linear Modulation)

As our final method on the complexity gradient, we look to modulate eat feature learned by the pLM and employ the established Feature-wise Linear Modulation [25] method to rescale assay-specific signals. Let $\mathbf{h} \in \mathbb{R}^{d_h}$ denote the frozen Stage 2 hidden representation (the output of the projection $\mathrm{Linear}(1280 \to d_h) + \mathrm{ReLU}$, with $d_h = 512$). The metadata embeddings and $\mathbf{h}$ are concatenated and passed through a two-layer MLP that produces per-sample modulation parameters:

$$[\boldsymbol{\gamma},\, \boldsymbol{\beta}] = \mathrm{MLP}_{\mathrm{FiLM}}\big([\mathbf{e}_a;\, \mathbf{e}_T;\, \mathbf{h}]\big) \in \mathbb{R}^{2d_h} \tag{8}$$

where $\mathrm{MLP}_{\mathrm{FiLM}}$ consists of $\mathrm{Linear}(d_a + d_T + d_h \to d_{\mathrm{film}})$, $\mathrm{ReLU}$, and $\mathrm{Linear}(d_{\mathrm{film}} \to 2d_h)$. The output is split into $\boldsymbol{\gamma}, \boldsymbol{\beta} \in \mathbb{R}^{d_h}$, and the modulated representation is:

$$\tilde{\mathbf{h}} = \boldsymbol{\gamma} \odot \mathbf{h} + \boldsymbol{\beta} \tag{9}$$

An optional post-modulation dropout is applied before the readout layer $\hat{y} = W_r\tilde{\mathbf{h}} + b_r$. A notable aspect of this design is that $\boldsymbol{\gamma}$ and $\boldsymbol{\beta}$ depend not only on the metadata $(\mathbf{e}_a, \mathbf{e}_T)$ but also on the sample representation $\mathbf{h}$. This allows the modulation to be input-dependent: the same assay label can induce different modulations for different pMHCs.

**Table 1:** Stage 3 conditioning architecture comparison.

| Variant | Trainable params | Conditioning mechanism | Init |
|---|---|---|---|
| Additive Fusion | $\sim$559K | $\mathbf{h} + \mathrm{MLP}([\mathbf{h}; \mathbf{e}_a; \mathbf{e}_T])$ | $\mathbf{r} = \mathbf{0}$ |
| FiLM | $\sim$810K† | $\boldsymbol{\gamma} \odot \mathbf{h} + \boldsymbol{\beta}$ | $\boldsymbol{\gamma}$=$\mathbf{1}$, $\boldsymbol{\beta}$=$\mathbf{0}$ |
| Calibration | $\sim$134K | $\alpha_g \hat{y}^{(S2)} + \beta_g + \Delta$ | $\alpha$=1, $\beta$=0, $\Delta$=0 |

† The production S3 model unfreezes the Stage 2 projection head increasing trainable parameters to $\sim$1.5M.

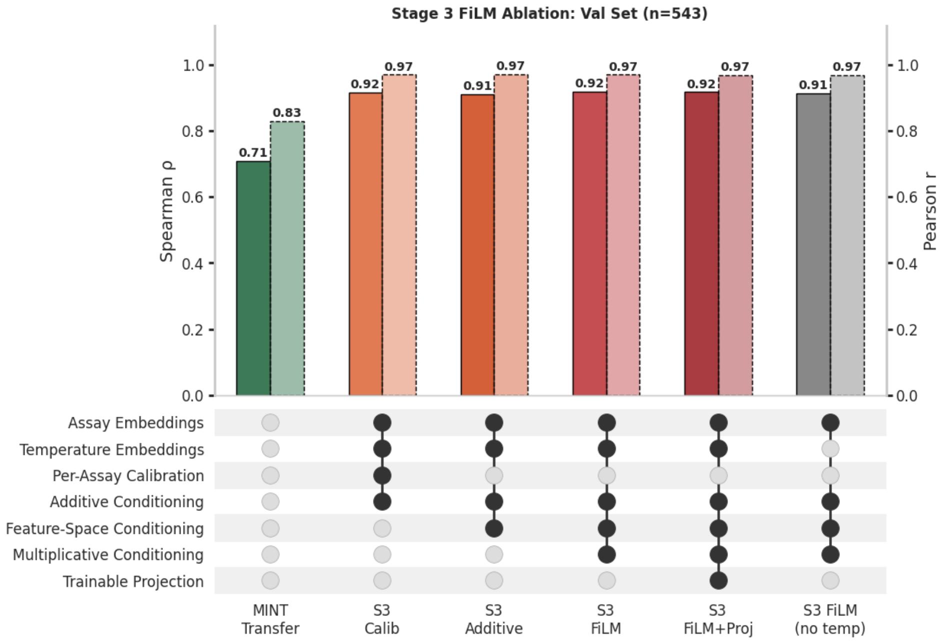

**Supplementary Figure 3:** *Assay Conditioning S3 Validation Performance.*

#### D Checkpoint Selection

Hyperparameters were selected via Bayesian optimization sweeps executed via Weights & Biases [26], with stage-specific objectives of minimizing validation loss for affinity (stage 1) and stability (stage 2) sweeps, and maximizing validation Spearman $\rho$ for the assay-conditioning (stage 3) sweeps given the mixed architectures and loss functions. Sweeps were run for MINT's two-stage training (individual sweeps for BA fine-tuning and BS fine-tuning), MINT's direct fine-tuning, and each of the three Stage 3 conditioning architectures. While analogous sweeps were conducted using ESM-2, the optimal hyperparameters did not vary substantially from those found for MINT, and as such the MINT architectural hyperparameters were applied to the ESM-2 baselines for a roughly parameter-equivalent comparison, while the optimizer and learning rate hyperparameters were left free to vary for a comparison between the maximally performant models. Later stage model sweeps were run on the optimal configurations of the previous stage (i.e. Stage 2 sweeps were run on the optimal Stage 1 checkpoints). For Stages 1 and 2, configs were run for 500 steps and the models with the lowest validation loss on the NetMHCstabpan partition were chosen; in the event of near ties, the configuration with the larger parameter count was selected, matched for learning rate. Stage 3 sweeps were run for with the same protocol but with Huber or MSE loss as options. These were run on the validation partition of the assay-conditioning data derived from the IEDB stability measurements for 500 steps as well. Table 2 summarizes the final hyperparameters for each model.

**Table 2:** Hyperparameters for ESM-2 and MINT stability prediction models.

| | **ESM-2** | | **MINT (S1/S2)** | | **Stage 3** | | |
|---|---|---|---|---|---|---|---|
| **Parameter** | Direct | Transfer | Direct | Transfer | Additive | Calib. | FiLM |
| Backbone | ESM2 | ESM2 | MINT | MINT | MINT | MINT | MINT |
| Cross-Attention | No | No | Yes | Yes | Yes | Yes | Yes |
| Learning Rate | $1{\cdot}10^{-4}$ | $1{\cdot}10^{-4}{\to}3{\cdot}10^{-5}$ | $1{\cdot}10^{-4}$ | $3{\cdot}10^{-5}{\to}3{\cdot}10^{-5}$ | $2.7{\cdot}10^{-5}$ | $9{\cdot}10^{-4}$ | $3{\cdot}10^{-5}$ |
| Hidden Dim | 256 | 512 | 256 | 512 | 512 | 512 | 512 |
| Dropout | 0.3 | 0.3 → 0.2 | 0.3 | 0.2 → 0.2 | 0.0 | 0.0 | 0.0 |
| Freeze % | 90% | 70%→70% | 90% | 50%→70% | 100% | 100% | 100%† |
| Batch Size | 16 | 16 | 16 | 16 | 16 | 16 | 16 |
| Loss | MSE | MSE | MSE | MSE | $\mathcal{H}(\delta$=1.0) | $\mathcal{H}(\delta$=0.5) | $\mathcal{H}(\delta$=1.0) |
| Assay Emb Dim | – | – | – | – | 32 | 4 | 32 |
| Temp Emb Dim | – | – | – | – | 32 | 4 | 8 |
| Cond. Hidden Dim | – | – | – | – | – | 256 | 512 |
| Trainable Params | – | – | – | – | ∼559K | ∼134K | ∼810† |

† The production S3 model unfreezes the Stage 2 projection head, increasing trainable parameters to ∼1.5M.

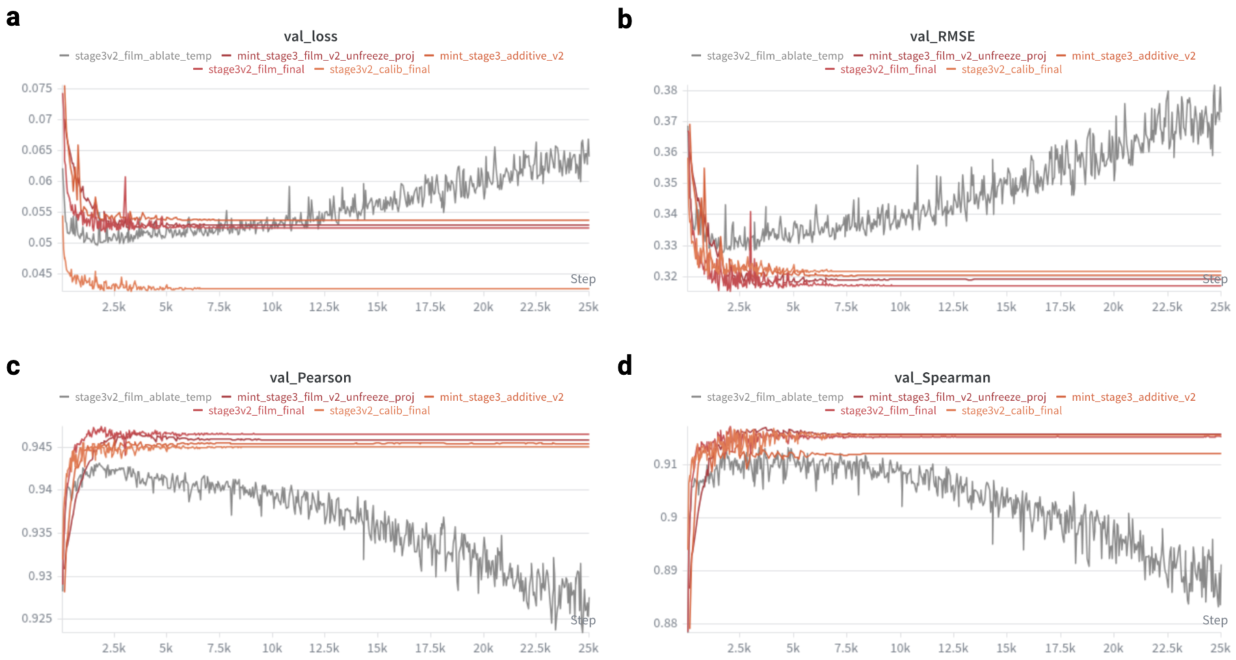

**Supplementary Figure 4:** *Assay Conditioning Training Curves.* **a.** Validation loss curves, **b.** RMSE, **c.** Pearson's $r$, and **d.** Spearman's $\rho$ for all Stage 3 models are shown across the entire training trajectory. Models include: affine residual calibration, additive latent fusion, FiLM, a FiLM variant with projection unfrozen, and a FiLM variant trained without temperature conditioning.

[3] James Robinson, Dominic J Barker, Xenia Georgiou, Michael A Cooper, Paul Flicek, and Steven GE Marsh. Ipd-imgt/hla database. *Nucleic acids research*, 48(D1):D948–D955, 2020.

[4] Michael Rasmussen, Emilio Fenoy, Mikkel Harndahl, Anne Bregnballe Kristensen, Ida Kallehauge Nielsen, Morten Nielsen, and Søren Buus. Pan-specific prediction of peptide–mhc class i complex stability, a correlate of t cell immunogenicity. *The Journal of Immunology*, 197(4): 1517–1524, 2016.

[5] Mikkel Harndahl, Michael Rasmussen, Gustav Roder, and Søren Buus. Real-time, high-throughput measurements of peptide–mhc-i dissociation using a scintillation proximity assay. *Journal of immunological methods*, 374(1-2):5–12, 2011.

[6] Randi Vita, Swapnil Mahajan, James A Overton, Sandeep Kumar Dhanda, Sheridan Martini, Jason R Cantrell, Daniel K Wheeler, Alessandro Sette, and Bjoern Peters. The immune epitope database (iedb): 2018 update. *Nucleic acids research*, 47(D1):D339–D343, 2019.

[7] Markus Wulf, Petra Hoehn, and Peter Trinder. Identification of human mhc class i binding peptides using the itopia™- epitope discovery system. In *Epitope Mapping Protocols: Second Edition*, pages 361–367. Springer, 2009.

[8] Arthur Fridman, Adam C Finnefrock, Daniela Peruzzi, Irene Pak, Nicola La Monica, Ansuman Bagchi, Danilo R Casimiro, Gennaro Ciliberto, and Luigi Aurisicchio. An efficient t-cell epitope discovery strategy using in silico prediction and the itopia assay platform. *Oncoimmunology*, 1 (8):1258–1270, 2012.

[9] Patrick A Ott, Zhuting Hu, Derin B Keskin, Sachet A Shukla, Jing Sun, David J Bozym, Wandi Zhang, Adrienne Luoma, Anita Giobbie-Hurder, Lauren Peter, Christina Chen, Oriol Olive, Todd A Carter, Shuqiang Li, David J Lieb, Thomas Eisenhaure, Evisa Gjini, Jonathan Stevens, William J Lane, Indu Javeri, Kaliappanadar Nellaiappan, Andres M Salazar, Heather Daley, Michael Seaman, Elizabeth I Buchbinder, Charles H Yoon, Maegan Harden, Niall Lennon, Stacey Gabriel, Scott J Rodig, Dan H Barouch, Jon C Aster, Gad Getz, Kai Wucherpfennig, Donna Neuberg, Jerome Ritz, Eric S Lander, Edward F Fritsch, Nir Hacohen, and Catherine J Wu. An immunogenic personal neoantigen vaccine for patients with melanoma. *Nature*, 547 (7662):217–221, July 2017.

[10] Zhuting Hu, Donna E Leet, Rosa L Allesøe, Giacomo Oliveira, Shuqiang Li, Adrienne M Luoma, Jinyan Liu, Juliet Forman, Teddy Huang, J Bryan Iorgulescu, Rebecca Holden, Siranush Sarkizova, Satyen H Gohil, Robert A Redd, Jing Sun, Liudmila Elagina, Anita Giobbie-Hurder, Wandi Zhang, Lauren Peter, Zoe Ciantra, Scott Rodig, Oriol Olive, Keerthi Shetty, Jason Pyrdol, Mohamed Uduman, Patrick C Lee, Pavan Bachireddy, Elizabeth I Buchbinder, Charles H Yoon, Donna Neuberg, Bradley L Pentelute, Nir Hacohen, Kenneth J Livak, Sachet A Shukla, Lars Rønn Olsen, Dan H Barouch, Kai W Wucherpfennig, Edward F Fritsch, Derin B Keskin, Catherine J Wu, and Patrick A Ott. Personal neoantigen vaccines induce persistent memory T cell responses and epitope spreading in patients with melanoma. *Nat. Med.*, 27(3):515–525, March 2021.

[11] David A Braun, Giorgia Moranzoni, Vipheaviny Chea, Bradley A McGregor, Eryn Blass, Chloe R Tu, Allison P Vanasse, Cleo Forman, Juliet Forman, Alexander B Afeyan, Nicholas R Schindler, Yiwen Liu, Shuqiang Li, Jackson Southard, Steven L Chang, Michelle S Hirsch,

Nicole R LeBoeuf, Oriol Olive, Ambica Mehndiratta, Haley Greenslade, Keerthi Shetty, Susan Klaeger, Siranush Sarkizova, Christina B Pedersen, Matthew Mossanen, Isabel Carulli, Anna Tarren, Joseph Duke-Cohan, Alexis A Howard, J Bryan Iorgulescu, Bohoon Shim, Jeremy M Simon, Sabina Signoretti, Jon C Aster, Liudmila Elagina, Steven A Carr, Ignaty Leshchiner, Gad Getz, Stacey Gabriel, Nir Hacohen, Lars R Olsen, Giacomo Oliveira, Donna S Neuberg, Kenneth J Livak, Sachet A Shukla, Edward F Fritsch, Catherine J Wu, Derin B Keskin, Patrick A Ott, and Toni K Choueiri. A neoantigen vaccine generates antitumour immunity in renal cell carcinoma. *Nature*, 639(8054):474–482, March 2025.

[12] Luis A Rojas, Zachary Sethna, Kevin C Soares, Cristina Olcese, Nan Pang, Erin Patterson, Jayon Lihm, Nicholas Ceglia, Pablo Guasp, Alexander Chu, Rebecca Yu, Adrienne Kaya Chandra, Theresa Waters, Jennifer Ruan, Masataka Amisaki, Abderezak Zebboudj, Zagaa Odgerel, George Payne, Evelyna Derhovanessian, Felicitas Müller, Ina Rhee, Mahesh Yadav, Anton Dobrin, Michel Sadelain, Marta Łuksza, Noah Cohen, Laura Tang, Olca Basturk, Mithat Gönen, Seth Katz, Richard Kinh Do, Andrew S Epstein, Parisa Momtaz, Wungki Park, Ryan Sugarman, Anna M Varghese, Elizabeth Won, Avni Desai, Alice C Wei, Michael I D'Angelica, T Peter Kingham, Ira Mellman, Taha Merghoub, Jedd D Wolchok, Ugur Sahin, Özlem Türeci, Benjamin D Greenbaum, William R Jarnagin, Jeffrey Drebin, Eileen M O'Reilly, and Vinod P Balachandran. Personalized RNA neoantigen vaccines stimulate T cells in pancreatic cancer. *Nature*, 618(7963):144–150, June 2023.

[13] Zachary Sethna, Pablo Guasp, Charlotte Reiche, Martina Milighetti, Nicholas Ceglia, Erin Patterson, Jayon Lihm, George Payne, Olga Lyudovyk, Luis A Rojas, Nan Pang, Akihiro Ohmoto, Masataka Amisaki, Abderezak Zebboudj, Zagaa Odgerel, Emmanuel M Bruno, Siqi Linsey Zhang, Charlotte Cheng, Yuval Elhanati, Evelyna Derhovanessian, Luisa Manning, Felicitas Müller, Ina Rhee, Mahesh Yadav, Taha Merghoub, Jedd D Wolchok, Olca Basturk, Mithat Gönen, Andrew S Epstein, Parisa Momtaz, Wungki Park, Ryan Sugarman, Anna M Varghese, Elizabeth Won, Avni Desai, Alice C Wei, Michael I D'Angelica, T Peter Kingham, Kevin C Soares, William R Jarnagin, Jeffrey Drebin, Eileen M O'Reilly, Ira Mellman, Ugur Sahin, Özlem Türeci, Benjamin D Greenbaum, and Vinod P Balachandran. RNA neoantigen vaccines prime long-lived CD8+ T cells in pancreatic cancer. *Nature*, 639(8056):1042–1051, March 2025.

[14] Ugur Sahin, Alexander Muik, Isabel Vogler, Evelyna Derhovanessian, Lena M. Kranz, Mathias Vormehr, Jasmin Quandt, Nicole Bidmon, Alexander Ulges, Alina Baum, Kristen E. Pascal, Daniel Maurus, Sebastian Brachtendorf, Verena Lörks, Julian Sikorski, Peter Koch, Rolf Hilker, Dirk Becker, Ann-Kathrin Eller, Jan Grützner, Manuel Tonigold, Carsten Boesler, Corinna Rosenbaum, Ludwig Heesen, Marie-Cristine Kühnle, Asaf Poran, Jesse Z. Dong, Ulrich Luxemburger, Alexandra Kemmer-Brück, David Langer, Martin Bexon, Stefanie Bolte, Tania Palanche, Armin Schultz, Sybille Baumann, Azita J. Mahiny, Gábor Boros, Jonas Reinholz, Gábor T. Szabó, Katalin Karikó, Pei-Yong Shi, Camila Fontes-Garfias, John L. Perez, Mark Cutler, David Cooper, Christos A. Kyratsous, Philip R. Dormitzer, Kathrin U. Jansen, and Özlem Türeci. Bnt162b2 vaccine induces neutralizing antibodies and poly-specific t cells in humans. *Nature*, 595(7868):572–577, Jul 2021. ISSN 1476-4687. doi: 10.1038/s41586-021-03653-6. URL `https://doi.org/10.1038/s41586-021-03653-6`.

[15] Jessica S W Borgers, Divya Lenkala, Victoria Kohler, Emily K Jackson, Matthijs D Linssen, Sebastian Hymson, Brian McCarthy, Elizabeth O'Reilly Cosgrove, Kristen N Balogh, Ekaterina Esaulova, Kimberly Starr, Yvonne Ware, Sebastian Klobuch, Tracey Sciuto, Xi Chen, Gauri Mahimkar, Joong Hyuk F Sheen, Suchitra Ramesh, Sofie Wilgenhof, Johannes V van Thienen, Karina C Scheiner, Inge Jedema, Michael Rooney, Jesse Z Dong, John R Srouji, Vikram R Juneja, Christina M Arieta, Bastiaan Nuijen, Claudia Gottstein, Olivia C Finney, Kelledy Manson, Cynthia M Nijenhuis, Richard B Gaynor, Mark DeMario, John B Haanen, and Marit M van Buuren. Personalized, autologous neoantigen-specific T cell therapy in metastatic melanoma: a phase 1 trial. *Nat. Med.*, 31(3):881–893, March 2025.
